# Fatty acid synthesis restricts HIV-1 infection through regulation of the nuclear envelope in CD4+ T cells

**DOI:** 10.64898/2026.09.22.753563

**Authors:** Joshua A. Acklin, Ming Ye, Erika A. Pontillo, Marc E. Wörös, Ethan H. Krupa, Bruce D. Walker, Alison E. Ringel

## Abstract

HIV-1 must breach intracellular defenses to establish infection in CD4+ T cells, yet the metabolic activities that sustain these barriers remain poorly understood. Using a metabolically targeted small molecule screen, we identify *de novo* fatty acid synthesis as an unexpected determinant of resistance to HIV-1 infection in activated CD4+ T cells. Inhibition of acetyl-CoA carboxylase 1 (ACC1) or fatty acid synthase (FASN) increased cellular susceptibility to infection, while fatty acid supplementation restored baseline levels of anti-viral resistance. Saturated fatty acids conferred the strongest protection, revealing that susceptibility depends on both cellular metabolic state and the composition of the extracellular lipid environment. Blocking fatty acid synthesis remodeled the cellular lipid pool, particularly for the membrane lipid phosphatidylcholine, and was accompanied by structural disruption of the nuclear envelope that serves as a physical barrier limiting passage of the HIV-1 capsid into the nucleus. FASN inhibition also selectively altered sensitivity to antiretroviral drugs that target the interaction between capsid and the nuclear pore. Broadly, these findings uncover a previously unrecognized role for cellular fat synthesis in the maintenance of the nuclear envelope and reveal that anti-viral pathways can emerge at the interface between cellular state and exposure.

## INTRODUCTION

HIV-1 depends on host cell machinery to establish productive infection. Once the virus integrates into the host genome, HIV-1 can persist in latently infected cells, requiring lifelong anti-retroviral suppression in most individuals. Defining cellular features that govern which cells become successfully infected is therefore critical to understanding the earliest events that form the latent HIV-1 reservoir, which comprises a major barrier to achieving a cure. Among cells that express the necessary co-receptors for viral entry, susceptibility to HIV-1 infection is heterogenous, with only a small subset of cells supporting productive infection^1,2^. Activation state is one factor that controls the susceptibility of CD4+ T cells to HIV-1 infection, as resting T cells are more refractory to infection compared to proliferating T cells^3^. However, the capacity to support infection varies even among activated T cells, and the underlying reasons are still incompletely understood^2,4^. Some of this variation can be explained by anti-viral restriction factors. For example, expressing APOBEC3 family members can restrict HIV-1 infection through the introduction of deleterious mutations into the viral genome, and interferon signals from neighboring cells can induce the expression of factors like MX2 by CD4+ T cells, preventing nuclear import of the viral genome^5–10^. However, the expression of these restriction factors does not completely explain why infection proceeds in some cells, but not others. This suggests that additional cellular factors exist which dictate susceptibility to HIV-1 infection.

HIV-1 also depends on metabolic support from the host cell. T cells with a high capacity for glucose uptake are more susceptible to HIV-1 infection, which is independent of activation or differentiation status^4,11^. Similarly, HIV-1 more readily infects T cells with greater mitochondrial mass, and redirecting glucose-derived carbon from lactate production toward mitochondrial oxidation enhances early viral reverse transcription^12^. Beyond supplying energy and biomolecules that HIV-1 can appropriate for constructing new viral particles, host metabolic activities maintain the organellar structures and signaling networks with which viral components must interact to establish productive infection. Thus, prior studies have identified central carbon metabolism as a positive regulator of HIV-1 infection, while raising the question of whether other metabolic pathways exert similar control.

Here, we sought to define metabolic pathways that modulate susceptibility to HIV-1 infection. To achieve this, we screened a curated small molecule library targeting cellular metabolism for compounds that alter the establishment of HIV-1 infection in primary human CD4+ T cells. Unexpectedly, most metabolic inhibitors increased the proportion of infected cells at concentrations that minimally affect viability, indicating that host metabolism largely restricts early stages of the HIV-1 lifecycle. Inhibitors of *de novo* fatty acid synthesis were highly represented among compounds that enhanced infection and induced a dependency on extracellular fatty acids to restore HIV-1 restriction. Mechanistically, inhibiting fatty acid synthesis altered the composition of membrane lipids, disrupted the bilayer architecture of the nuclear envelope, and increased the potency of anti-retroviral drugs that target the nuclear import of HIV-1 capsid. These findings identify *de novo* fatty acid synthesis as an environmentally responsive HIV-1 restriction pathway and demonstrate that viral susceptibility can emerge from the interplay between host cell biology and the extracellular environment.

## RESULTS

### Small molecule screening identifies metabolic states of HIV-1 susceptibility

Given that HIV-1 requires host metabolic resources to support its viral lifecycle and navigates cellular structures composed of host-derived biomolecules, we reasoned that metabolic pathways may serve as restriction or dependency factors. We sought to examine this possibility systematically by evaluating the impact of a metabolically targeted small-molecule screen on HIV-1 infection efficiency in primary human CD4+ T cells^13^. Because T cell activation increases susceptibility to HIV-1 and simultaneously elicits extensive metabolic remodeling^3,14^, we designed the screen to minimize identifying compounds that modulate infection via activation. Primary human CD4+ T cells, procured from StemCell Technologies from donors with similar demographics and CD4+ T cell subset ratios (**Figure S1A-B**), were expanded with anti-CD3/CD28 dynabeads in the presence of recombinant human IL-2 to generate a pool of antigen-experienced cells in sufficient numbers to perform the entire screen. To perform the screen, T cells were acutely stimulated for 24 hours in bulk and then distributed across 384-well plates for pre-treatment with the small-molecule library for another 24 hours, which consisted of 240 compounds across a 10-point titration curve (**Figure 1A**). Importantly, the activation signal was limited to the stimulation phase to separate the effect of metabolic perturbation from cell activation. This design also provided time for the T cells to adapt to each metabolic perturbation before interacting with HIV-1. Cells were then infected with a single-cycle VSV-G pseudotyped HIV-1 reporter virus (NL43 ΔENV GFP-VSVG), and viral susceptibility was measured 72 hours later by high-throughput flow cytometry as the proportion of infected cells based on expression of the GFP reporter. This enabled the effect of metabolic perturbations to be examined on a single round of the viral lifecycle, as pseudotyped HIV-1 cannot generate new infectious particles.

**Figure 1:**
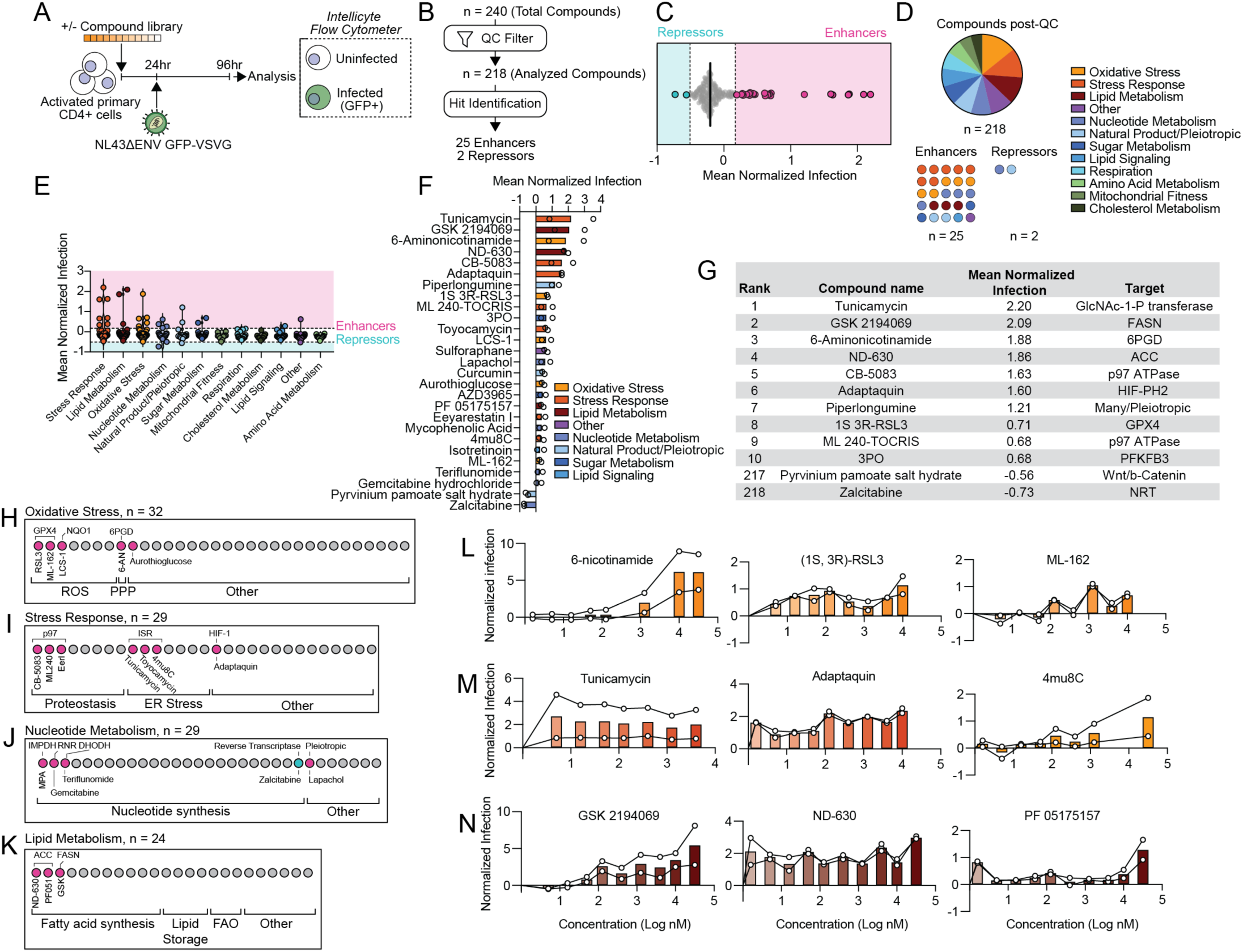
Small molecule screening identifies metabolic states of HIV-1 susceptibility. (A) Schematic of experimental design of high-throughput infection screening. Primary CD4+ T cells isolated from healthy human donors were activated for 24 hours and then treated with 240 inhibitors. After 24 hours of inhibitor treatment, cells were infected with VSVG pseudotyped HIV-1 (VSVG-NL43 ENV-GFP) for 72 hours. Infection was read out as %GFP+ cells. The screen was performed with two biological replicates (i.e. cells from two healthy human donors). (B) Overview of data filtering and hit identification pipeline. (C) Mean normalized infection of all compounds remaining after quality control filters, highlighting enhancers and repressors. Dashed line indicates a Tukey’s fence (k=1) which was utilized to define enhancers and repressors. (D) Organization of compounds into 12 categories based on cellular targets. (E) Mean normalized infection for compounds in each category. (F) Rank-ordered bar plot of mean normalized infection of all enhancers and repressors. Datapoints represent mean normalized infection value for each donor (n = 2). Bars represent average of two donors. (G) Summary table of mean normalized infection and cellular targets of top 10 enhancers and all repressors. (H) Breakdown of enhancers in the oxidative stress category. Each circle represents a compound in the category, with compound targets annotated above the circles and compound names below. Pink circles indicate enhancers, while blue indicate repressors. (I) As with (H), but with enhancers in stress response category. (J) As with (H), but with enhancers and repressors in nucleotide metabolism category. (K) As with (H), but with enhancers in the lipid metabolism category. (L) Titration curves of highlighted enhancers in oxidative stress category. Individual titration curves represent normalized infection value for each donor. (M) As with (L), but with enhancers in stress response category. (N) As with (L), but with enhancers in lipid metabolism category. *Abbreviations:* QC = Quality Control; FASN = Fatty acid synthase; 6PGD = 6-phosphogluconate dehydrogenase; ACC = Acetyl-coA Carboxylase; HIF-PH2 = Hypoxia-inducible factor prolyl hydroxylase 2; GPX4 = Glutathione Peroxidase 4; PFKFB3 = 6-phosphofructo-2-kinase/fructose-2,6-biphosphatase 3; ROS = reactive oxygen species; PPP = pentose phosphate pathway; ISR = integrated stress response; ER-endoplasmic reticulum; FAO = fatty acid oxidation. NQ01 = NAD(P)H quinone dehydrogenase 1; IMPDH = Inosine-5′-monophosphate dehydrogenase; RNR = ribonucleotide reductase; DHODH = dihydroorotate dehydrogenase.

To examine the impact of metabolic perturbations on HIV-1 infection, we first defined a normalized infection metric that corrected for plate-to-plate variation. With this metric, compounds that do not impact infection have a value of 0, while compounds that increase or reduce infectious susceptibility have positive or negative scores, respectively, with full repression of infection represented as -1 (**Fig. S1C**). Each plate contained vehicle and Raltegravir controls for normalization. Raltegravir is a potent integrase inhibitor used to establish the lower bound for infection^15,16^. After scaling raw viability and infection percentages to vehicle and Raltegravir treatment on a per-plate basis, compounds were removed that failed to pass a quality control filter (**Figure S1C-D**). Among the 218 of 240 compounds remaining, we applied a secondary filter to retain compounds with at least three concentrations that 1) maintained >= 90% viability compared to vehicle control and 2) displayed concordant infection outcomes across both donors (**Figure 1B, Figures S1E-F**). We next applied a modified Tukey’s fence (k = 1) on mean normalized infection across all doses to identify screen hits above a minimum effect size. This analysis yielded 25 enhancers and 2 repressors (**Figure 1C**). The enhancers largely belonged to stress response (n=7), oxidative stress (n=5), nucleotide metabolism (n=4), and lipid metabolism categories (n=3) (**Figures 1D-E**). The top repressor identified was a nucleoside analogue called Zalcitabine that inhibits reverse transcriptase^17^ (**Figure 1F-G**). As reverse transcription of viral genetic material is required prior to integration into the host genome, this provides further validation that our screen and analytical pipeline reports on early HIV-1 infection.

Because most of the hits that emerged from this screen enhanced rather than repressed infection, we examined whether common cellular pathways were represented among the 25 enhancer compounds. We initially focused on the four metabolic categories with the highest mean normalized infection (**Figure 1E**, **1H-K**). Among the 32 compounds assigned to the oxidative stress category, 5 enhanced HIV-1 infection (**Figure 1H**). Two of these compounds targeted glutathione peroxidase 4 (GPX4), which uses reduced glutathione to detoxify lipid peroxides that cause ferroptosis (**Figures 1H, 1L**)^18^. The pentose phosphate pathway inhibitor 6-aminonicotinamide (6-AN) also strongly enhanced infection (**Figures 1H, 1L**). 6-AN inhibits 6-phosphogluconate dehydrogenase (6PGD), an enzyme that maintains the high NADPH to NADP+ ratio needed for reducing the cellular glutathione pool as well as performing reductive biosynthetic reactions^19^. Of the 29 compounds assigned to the stress response category, 7 enhanced HIV-1 infection (**Figure 1I**). Three of these compounds (CB-5083, ML240, and Eeyarestatin I) inhibit the p97 ATPase, which protects against proteotoxic stress through several mechanisms, including extracting misfolded proteins from the endoplasmic reticulum (ER) (**Figure 1I**)^20–22^. Three other hits in this category also perturb ER homeostasis, including tunicamycin, which causes ER protein misfolding, as well as toyocamycin, and 4mu8C that inhibit the resulting unfolded protein response (**Figures 1I, 1M**). Adaptaquin, an inhibitor of HIF prolyl hydroxylases, also enhanced infection across nearly all concentrations tested (**Figures 1I, 1M**). The screen also identified multiple hits in the nucleotide metabolism category that enhance infection, all of which are involved in nucleotide synthesis. Inhibiting dihydroorotate dehydrogenase (DHODH), inosine-monophosphate dehydrogenase (IMPDH), and ribonucleotide reductase (RNR) all enhanced the establishment of infection in this assay (**Figure 1J**). It is important to note that gemcitabine also has reported anti-retroviral activity^23^. However, the anti-viral effect of gemcitabine treatment occurs outside the early infection window, as it decreases the fitness of viral progeny by mutagenizing the viral genome, and therefore would not be apparent in this screen^23^. Lastly, we considered the 24 compounds in the lipid metabolism category. All 3 hits in this category were inhibitors of fatty acid synthesis (**Figures 1K, 1N**). Both ND-630 and PF-05175157 inhibit acetyl-CoA carboxylase (ACC), which catalyzes the conversion of acetyl-CoA to malonyl-CoA for fatty acid synthesis^24,25^. GSK2194069 targets FASN, which uses acetyl-CoA and malonyl-CoA to synthesize long-chain fatty acids. This connects to the identification of 6-AN as an enhancer from the oxidative stress category, which may also restrict fatty acid synthesis by limiting the NADPH required for each round of elongation catalyzed by FASN (**Figures 1H**). Taken together, these results reveal multiple metabolic pathways that control permissiveness to HIV-1 infection and nominate fatty acid synthesis as a key control point on HIV-1 susceptibility.

### Fatty acid Synthesis restricts HIV-1 infection in CD4+ T cells

The small molecule screen presented in **Figure 1** revealed that multiple steps in *de novo* fatty acid synthesis restricted HIV-1 infectivity. Fatty acid synthesis is a stepwise pathway that uses citrate as a carbon source to build palmitate through iterative cycles of carbon addition^26,27^. Cytosolic citrate is first converted into acetyl-CoA by the enzyme ATP-citrate lyase (ACLY), which is subsequently carboxylated to malonyl-CoA by the enzyme acetyl-CoA carboxylase (ACC). Acetyl-CoA and malonyl-CoA are then used by FASN to build an acyl chain through repeated condensation reactions, which produces a fully saturated 16-carbon fatty acid called palmitate that can be further elongated or desaturated to generate the cellular fatty acid pool (schema in **Figure 2A**) ^reviewed in^ ^28^. We independently validated that the ACC inhibitor (ND630)^25^ and FASN inhibitor (GSK2194069)^29^ in the small molecule screen sensitized CEM-GXR cells, a CD4+ T cell cancer cell line used to study HIV-1 virology^30^, to HIV-1 infection. We observed a dose-dependent increase in infection for both compounds (**Figures 2B-C**), consistent with the screen outcome. While the ACC inhibitor had no impact on viability at any dose tested, inhibiting FASN induced a 15% decrease in viability at the maximal dose of inhibitor that did not achieve statistical significance (**Figure S2A-B**). Both inhibitors blocked fatty acid synthesis in T cells at concentrations that raised HIV-1 infectivity, as measured by stable isotope tracing. While CEM-GXR cells cultured in uniformly labeled ^13^C_6_-glucose incorporated glucose-derived carbon into palmitate, treatment with either ND630 or GSK2194069 completely abrogated these labeling patterns (**Figure 2D-E**). Lastly, we corroborated the effect of FASN inhibition on HIV-1 infection using a structurally distinct inhibitor, TVB2640^31^, which was not present in the small-molecule screen. TVB2640 treatment also caused a dose-dependent increase in infectivity for CEM-GXR cells (**Figure 2F**). Like GSK219469, there was a significant but minor decrease in viability compared to DMSO-treated controls (∼15%; **Figure S2C**). As FASN acts downstream of ACC1, we focused on inhibitors of FASN for further analysis.

**Figure 2.**
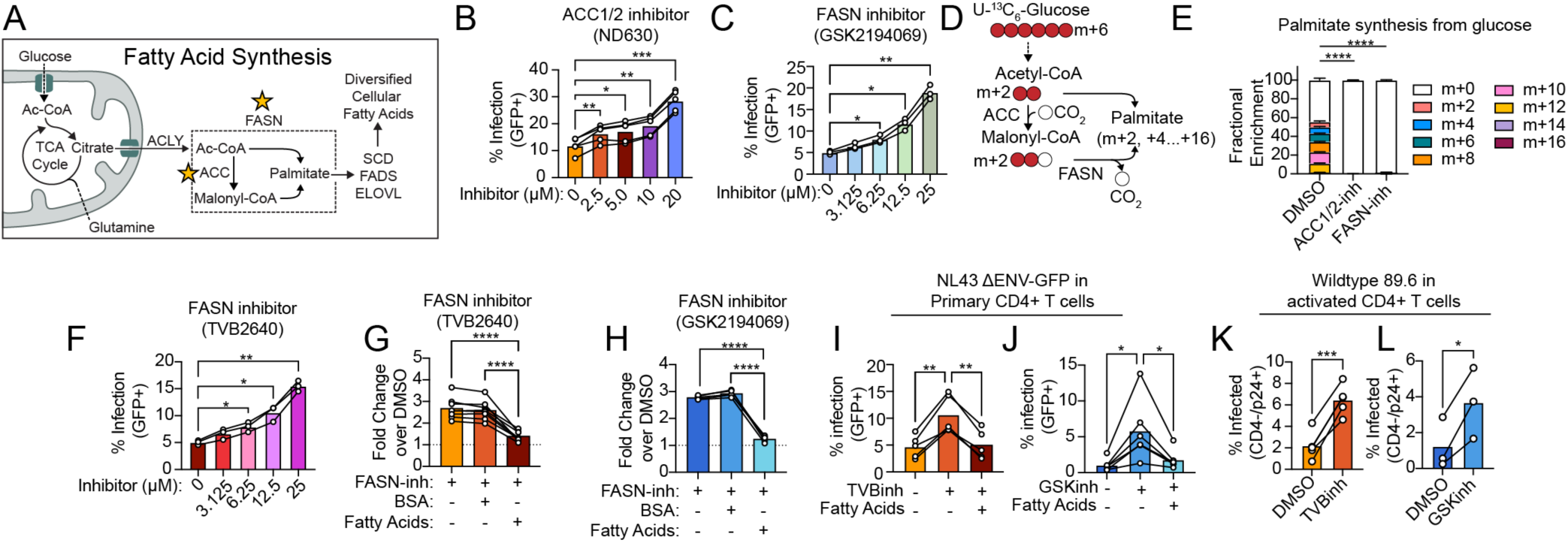
Inhibition of fatty acid synthesis enhances infectivity of HIV-1. The potential for cellular fatty acid synthesis to regulate HIV-1 infectivity was tested across a variety of infection systems. Unless denoted otherwise, each datapoint represents an experimental replicate, which is the average of three technical replicates. (A) Overview of fatty acid synthesis in human T cells. Palmitate, a 16-carbon saturated fatty acid from which the diversified cellular fatty acid pool is derived, is synthesized from TCA-cycle derived citrate in a series of sequential enzymatic reactions in the cytosol. Enzymes targeted in this figure (ACC and FASN) are starred. The dashed box encompasses the FASN reaction, which utilizes the substrate and product of the ACC1 step. (B) Impact of ACC inhibitor ND630 (identified in Figure 1) on infectivity of VSV-G pseudotyped HIV-1 reporter virus (NL43ΔENV GFP-VSVG) in CEM-GXR cells (N=4 experimental replicates). Statistical significance was determined by one-way ANOVA with Dunnet’s multiple comparison test against the drug-control. (C) As with (B), but with FASN inhibitor GSK2194069 (N=3 experimental replicates). (D) Overview of stable isotope tracing of uniformly labeled ^13^C glucose into palmitate synthesis. The extent of isotopic labeling is denoted as m+ the number of ^13^C carbons incorporated in each molecule. (E) Stable isotope tracing of uniformly labeled ^13^C glucose under DMSO, ACC inhibition (ND630; 25μM) or FASN inhibition (GSK2194069; 25μM). Palmitate was derivatized into palmitate methyl-ester, and fractional enrichment of labeled carbon in palmitate methyl-ester is shown, indicating relative activity of cellular fat synthesis in each condition (N=3 technical replicates per group). Statistical significance is only shown for a comparison of the no-label fraction in each group. DMSO control maintained a p-value ≤ 0.05 for all measured isotopologues against ND630 and GSK-treated conditions. (F) As with (B-C), but with an additional FASN inhibitor, TVB2640, that was not included in the small molecule screen from Figure 1 (N=3 experimental replicates). (G) Capacity of exogenous fatty acids (palmitate-BSA and oleate-BSA; each 25μM) to rescue HIV-1 susceptible state generated with FASN-inhibitor TVB2640 as compared to fatty-acid free BSA, which is the vehicle for exogenous fatty acids (N=8 experimental replicates). Statistical significance was determined by one-way ANOVA with Tukey’s multiple comparison test. (H) As with (F), but with FASN-inhibitor GSK2194069 (N=4 experimental replicates). (I) Impact of FASN inhibitor TVB2640 and exogenous fatty acid add-back (25μM) on infectivity of NL43ΔENV-GFP in acutely activated primary CD4+ T cells (N=5 donors). Statistical significance was determined by one-way ANOVA with Tukey’s multiple comparison test. (J) As with (H), but with FASN inhibitor GSK2194069 (N=5 donors). (K) Impact of FASN inhibitor TVB2640 (25μM) on infectivity of replication competent HIV^89.6^ in acutely activated primary CD4+ T cells (N=4 donors). Statistical significance was determined by paired t-test. (L) As with (J) but with FASN inhibitor GSK2194069 (N=3 donors). *Statistical Significance:* *p<0.05, **p≤0.01, ***p≤0.001, ****p≤0.0001. *Abbreviations*: Ac-CoA = Acetyl-coA; ACLY = ATP citrate lyase; ACC = Acetyl-coA carboxylase; FASN = Fatty acid synthase; SCD = stearoyl-CoA desaturase 1; FADS2 = Fatty acid desaturase 2; ELOVL = Elongation of very long-chain fatty acids protein; m = degree of isotopic labeling; FASNinh = FASN inhibitor; BSA = Bovine serum albumin (fatty-acid free); DMSO = dimethyl sulfoxide.

FASN inhibition could increase infectivity by depleting its product palmitate or by making its substrates (NADPH, malonyl-CoA, and acetyl-CoA) available for other cellular processes. To distinguish between these possibilities, we evaluated whether palmitate supplementation prevented the increase in infectivity with FASN inhibition. We delivered palmitate conjugated to bovine serum albumin (BSA) to mimic how non-esterified fatty acids circulate in the bloodstream and included fatty acid-free BSA as a negative control. To avoid palmitate toxicity that has been previously noted in T cells^32,33^, we delivered palmitate-BSA in combination with oleate-BSA at equimolar concentrations, which will subsequently be referred to as “exogenous fatty acids”. As in previous experiments, both TVB2640 (**Figure 2G**) and GSK2194069 (**Figure 2H**) enhanced infectivity in CEM-GXR cells over DMSO-treated controls. Exogenous fatty acids were sufficient to prevent this increase in infectivity for both compounds, while fatty-acid-free BSA did not impact the HIV-1 susceptible state (**Figure 2G-H**). Similarly, exogenous fatty acids restored viability due to FASN inhibition (**Figure S2D**). This effect was conserved in acutely activated primary CD4+ T cells from healthy human donors, where FASN inhibition likewise increased susceptibility to HIV-1 infection in a manner that could be prevented by exogenous fatty acids (**Figure 2I-J, Figure S2E-F**). FASN inhibition also reduced primary T cell proliferation in a manner that could be restored by fatty acids, as demonstrated by retention of CellTrace Violet (CTV) dye (**Figure S2G**). However, proliferation was similarly restricted in both infected and uninfected cells from the same culture, indicating that changes in susceptibility are not due to a greater expansion of infected cells over uninfected counterparts (**Figure S2H**). We confirmed that exogenous fatty acids could be taken up by CD4+ T cells by measuring uptake of a click-chemistry compatible palmitate analogue (palmitate-azide) after reacting it with alkyne-modified Alexa Fluor 647 dye (**Figure S2I**)^34–36^. Primary T cells took up palmitate-azide readily at 37°C across all treatment groups relative to cold temperatures that slow transport (**Figure S2J**). Together, these findings indicate that the depletion of intracellular palmitate during FASN inhibition reduces viability and proliferation and drives an increase in susceptibility to HIV-1 infection.

Finally, we extended our analysis to infection by the replication-competent HIV-1 dual-tropic isolate 89.6 in primary CD4+ T cells, which enters host cells through CD4 and CCR5/CXCR4^37^. This enabled us to exclude the possibility that increased susceptibility during FASN inhibition was mediated by the artificial entry route utilized by VSV-G pseudotyped viruses^38^. Even in the context of replication-competent, wildtype HIV-1 entry, FASN inhibitors increased infectivity (**Figure 2K-L**). Taken together, these data demonstrate a role for cellular fat synthesis in restricting HIV1 infection in a manner that depends upon the capacity to acquire palmitate.

### Fatty acid synthesis shapes the properties of membrane lipids in CD4+ T cells

To nominate potential mechanisms by which fatty acid synthesis regulates HIV-1 infectivity, we conducted quantitative shotgun lipidomics on CEM-GXR cells that had been treated with FASN inhibitor (GSK2194069) plus fatty acid-free BSA, FASN inhibitor with exogenous fatty acids (palmitate-BSA + oleate-BSA), or DMSO plus fatty acid-free BSA as a vehicle control. At a high level, FASN inhibition induced a global reduction in total lipid abundance in T cells compared to vehicle-treated controls, while exogenous fatty acids increased total lipid content (**Figure 3A**). These sum abundances consisted of a total of 868 unique lipid species found in mammalian cells, spanning neutral lipids, phospholipids, sterols and sphingolipids. FASN inhibition caused the largest magnitude changes to neutral lipids (**Figure S3A**), prompting us to evaluate whether building or breaking down lipid droplets that store neutral lipids modulates HIV-1 infectivity. We tested two different inhibitors of diglyceride acyltransferase (DGAT1), the enzyme that catalyzes the final step in the synthesis of triacylglycerides^39^, which led to a modest decrease in HIV-1 infectivity (**Figure S3B-C**). Thus, inhibiting DGAT1 does not phenocopy the effect of FASN inhibition state or interfere with the impact of adding exogenous fatty acids to FASN-inhibited cells (**Figure S3B-C**).

**Figure 3:**
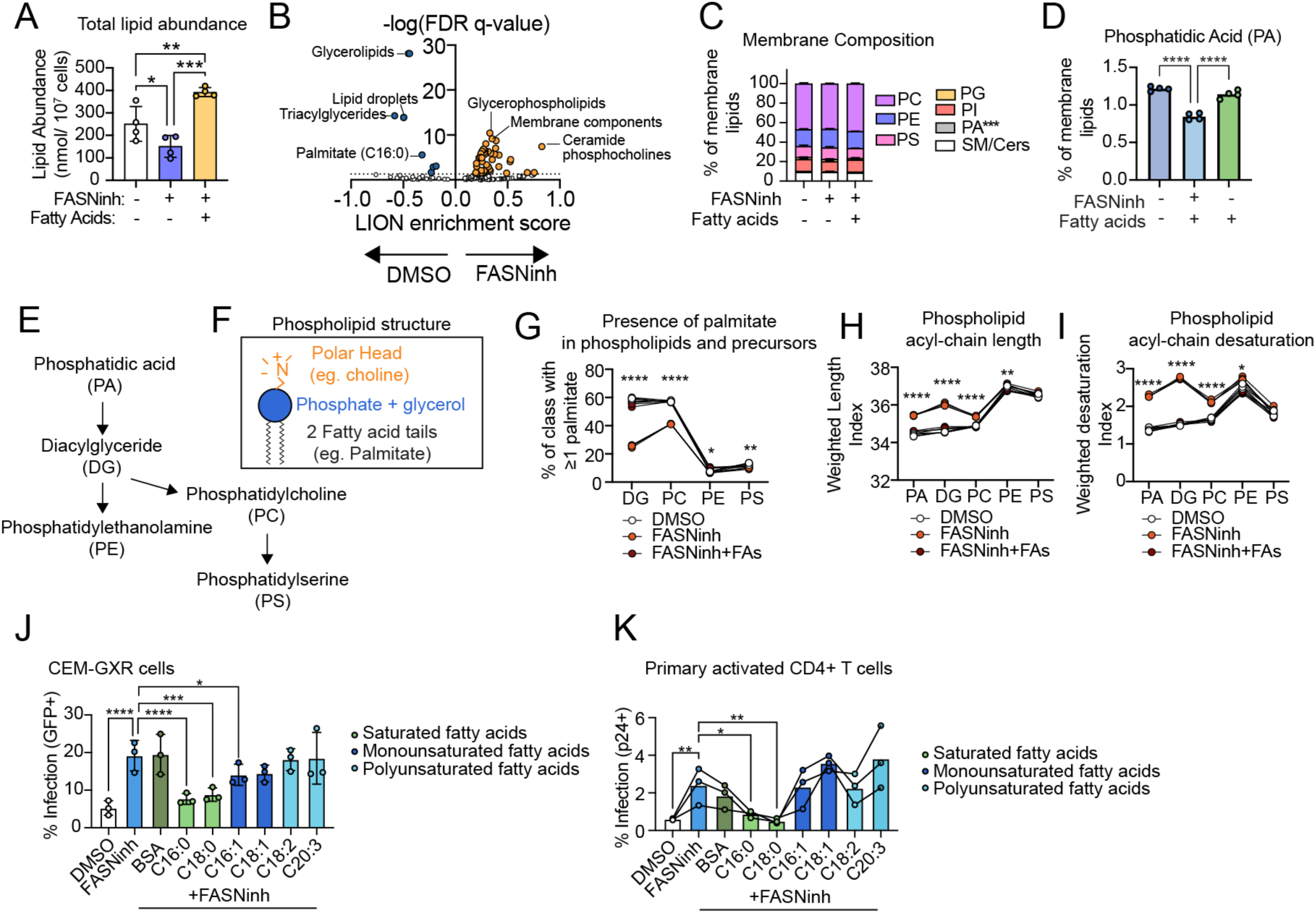
FASN inhibition remodels membrane phospholipids. (A) Comparison of total abundance for all measured lipid species across treatment group. Statistical significance was determined by one-way ANOVA with Holm-Sidak’s multiple comparison test. (B) Enrichment analysis of hierarchical lipid ontologies in FASN-inhibited condition as compared to DMSO-vehicle controls, as calculated utilizing the LION-web based ontology for lipid pathway enrichment^40^. Hits that maintained an FDR-adjusted q-value of ≤ 0.05 were indicated with orange (FASN-inhibited) or blue (DMSO). (C) The composition of membrane lipids is quantified and compared across groups as a percentage of total membrane lipid. Statistical significance was determined by one-way ANOVA with Tukey’s multiple comparison test was conducted within each class. (D) Comparison of the abundance of phosphatidic acid (PA) between treatment groups. Statistical significance was determined by one-way ANOVA with Tukey’s post-hoc test for multiple comparisons. (E) Overview of phospholipid synthesis from phosphatidic acid and diacylglycerol, which donate two fatty acid tails and a glycerol to the synthesis of major phospholipids phosphatidylethanolamine (PE), phosphatidylcholine (PC) and phosphatidylserine (PS).(F) General structure of a phospholipid, including a polar head group, phosphate/glycerol backbone, and two fatty acid tails. (G) The percentage of each major membrane lipid class (or precursor) that has at least one palmitate tail is assessed and compared between treatment conditions. Statistical significance is calculated by one-way ANOVA with Tukey’s post-hoc test for multiple comparisons within each subclass. (H-I) Comparison of the average sum length (H) or desaturation (I) of the fatty-acid tails in each of the major phospholipid subclasses and precursors across treatment. Statistical significance was determined by one-way ANOVA with Tukey’s multiple comparison test was conducted within each class. Stars indicate comparison between the FASN-inhibitor and DMSO-treated controls. (J-K) Potential for BSA-conjugated fatty acids of increasing length and desaturation to prevent enhanced HIV-1 infectivity induced by FASN inhibition at equivalent concentration in CEM-GXR cells (J) or primary human CD4+ T cells (K). Each data point represents an experimental replicate, which is the average of three technical replicates. Statistical significance was determined by Two-way ANOVA with Dunnet’s multiple comparison test against the FASN-inhibited condition. *Statistical Significance:* *p<0.05, **p≤0.01, ***p≤0.001, ****p≤0.0001. *Abbreviations: FASNinh =* FASN inhibitor (GSK2194069); LION = Lipid Ontology; DMSO = dimethyl sulfoxide; PC = phosphatidylcholine; PE = phosphatidylethanolamine; PS = phosphatidylserine; PG = phosphatidylglycerol; PI = phosphatidylinositol; PA = phosphatidic acid; SM = sphingomyelin; Cers = ceramides; DG = diacylglycerol.

To uncover biologically meaningful patterns in the lipidomic response to FASN inhibition, we analyzed the data using LION, a lipid ontology enrichment tool that groups lipids into hierarchical sets by shared structural properties and participation in cellular pathways^40^. We first compared the control condition (DMSO treatment) to CEM-GXR cells treated with FASN inhibitor. Consistent with the role of FASN in synthesizing palmitate, we found a statistically significant reduction in the ontology term for C16:0 palmitate in FASN-inhibited cells as compared to DMSO-vehicle treated controls (**Figure 3B**). FASN inhibition reduced the enrichment of glycolipid- and lipid droplet-associated categories, while enriching for membrane lipid categories, including glycerophospholipids, membrane components, and ceramide phosphocholines (**Figure 3B**). Treatment with exogenous fatty acids prevented these lipidomic shifts (**Figure S3D**), indicating that exogenous fatty acids could bypass lipidomic changes caused by loss of *de novo* fatty acid synthesis.

The results from LION analysis prompted us to hypothesize that alterations in membrane lipids were responsible for the increased infectivity associated with FASN inhibition. However, membrane lipid composition was remarkably stable across treatment conditions at a subclass level (**Figure 3C**), with the exception of phosphatidic acid (PA), which was selectively depleted by FASN inhibition (**Figure 3D**). PA is a precursor for glycerophospholipid synthesis (**Figure 3E-F**) that is also present at sites of negative membrane curvature^41–43^ and has been implicated in maintaining the architecture of the nuclear envelope in yeast^44,45^. In contrast, FASN inhibition caused pronounced changes in the acyl chain composition within diacylglyceride (DG) and major phospholipid subclasses, characterized by a global depletion of palmitate in the phospholipid pool that was particularly apparent for DG and phosphatidylcholine (PC) (**Figure 3G**). To compensate, we observed a concomitant increase in both the average length (**Figure 3H**) and number of unsaturated sites (**Figure 3I**) on fatty acyl chains that comprise the phospholipid pool during FASN inhibition. Providing exogenous fatty acids concurrently with the FASN inhibitor prevented these changes in chain length and unsaturation (**Figure 3H-I**). As with palmitate depletion, this was particularly striking in the precursor species (PA and DG) as well as within PC (**Figure 3H-I**). If this acyl-chain remodeling drives increased HIV-1 susceptibility, then only fatty acids with structural properties similar to palmitate should prevent the susceptible state induced by FASN inhibition. To test this, we pre-treated CEM-GXR cells with FASN inhibitor together with BSA-conjugated fatty acids of increasing length and saturation before challenging the cells with HIV-1 and quantifying infectivity. As shown in **Figure 3J**, only palmitate (C16:0), stearate (C18:0) and palmitoleic acid (C16:1) could significantly prevent the HIV-1 susceptible state, while longer, more unsaturated fatty acids had no impact. Similarly, only palmitate and stearate prevented the loss of viability caused by FASN inhibition (**Figure S3E**).

We next repeated these experiments in acutely activated primary CD4+ T cells to determine whether this pattern of susceptibility extended to primary cells. In the absence of FASN inhibition, individual fatty acids produced significant but small effects on HIV-1 infectivity as compared to the increase caused by the FASN inhibitor (**Figure S3F**). Similar to CEM-GXR cells, both palmitate (C16:0) and stearate (C18:0) returned primary T cell infectivity to baseline levels while monounsaturated and polyunsaturated fatty acids had no impact on the HIV-1 susceptible state (**Figure 3K**). There were minimal effects on viability for all fatty acids tested (**Figure S3G-S3H**). In sum, these data indicate that FASN inhibition alters the fatty acid composition of membrane phospholipids and suggest that the shift toward longer and more unsaturated fatty acids in membrane lipids may enhances HIV-1 infectivity.

### Fatty acid synthesis contributes to HIV-1 restriction by impacting nuclear translocation of capsid

To access host chromatin for integration, the HIV-1 capsid must translocate through the nuclear pore complex largely intact, making nuclear entry an important barrier to infection^46,47^. The nucleus is enclosed by two lipid bilayers that connect at nuclear pore complexes^48–50^, prompting us to examine whether lipidomic remodeling induced by FASN inhibition impacted the integrity of this structure critical to the viral lifecycle. Transmission electron microscopy (TEM) in CEM-GXR cells revealed striking changes in the nuclear envelope morphology associated with FASN inhibition. In control cells, the inner and outer membranes of the nuclear envelope were distinct and separated by a narrow perinuclear space (**Figure 4A**). Under FASN inhibition, the nuclear envelope became highly irregular. The coordination of the inner and outer nuclear membranes was disjointed, resulting in perinuclear widening (**Figure 4B**). With exogenous fatty acid treatment, the nuclear envelope normalized, returning to the baseline state with occasional, but smaller, areas of perinuclear widening (**Figure 4C**). These changes were limited to the nuclear envelope, as cells treated with FASN inhibitor were otherwise unremarkable (**Figure S4A**) and of similar total size (**Figure S4B**).

**Figure 4:**
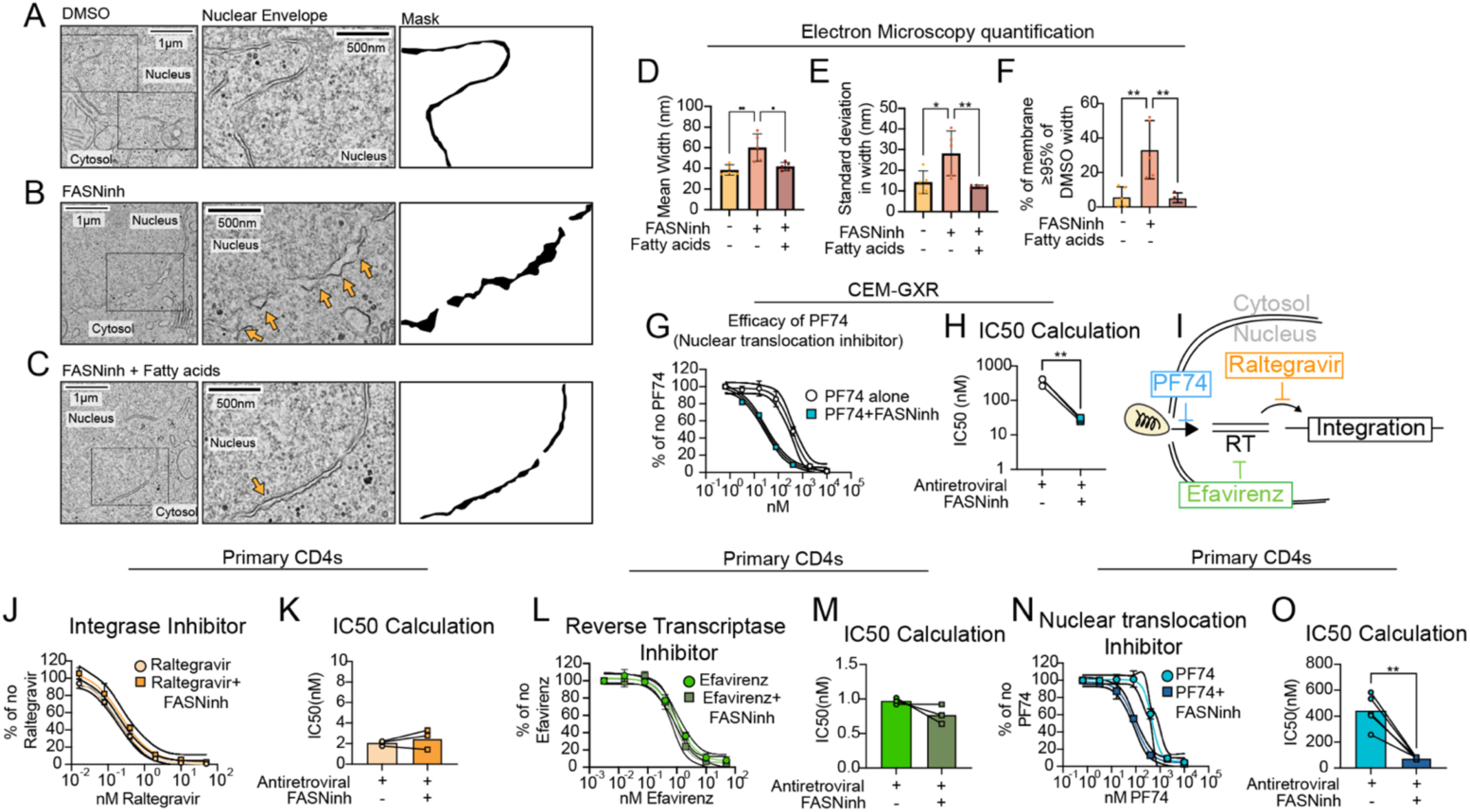
FASN inhibition enhances the efficacy of antiretroviral targeting nuclear translocation of capsid. (A-C) Representative 11,000x image of the nuclear/cytoplasmic border is shown (left) with a cropped zoom of the nuclear envelope in DMSO treated controls (A), FASN-inhibited cells (B), or in FASN-inhibited cells with exogenous palmitate-BSA and oleate-BSA (C). Scale bars denote 1 µm lengths in left panels, or 500 nm lengths in middle panels. A manually traced mask of the nuclear envelope in each condition is shown on the right. (D) Quantification of the mean width of the perinuclear space and comparison between groups. 5 cells per group were manually segmented in FIJI, and the average width between the outer and inner nuclear envelope were quantified every 2 nm. Each data point represents the average of 4-6 segments quantified from each image. Significance was determined by one-way ANOVA with Tukey’s post-hoc test for multiple comparisons. See *Methods*. (E) As with (D) but quantification of the standard deviation in the width of each measured nuclear envelope segment. (F) The 95^th^ percentile was calculated for each nuclear envelope segment and averaged per cell in the DMSO controls. The percentage of measured widths equal to or exceeding this value represent the percentage of nuclear envelope considered “expanded” in each group. Significance was determined as in (D-E). (G) The sensitivity of infection to antiretroviral PF74 was examined in cells pre-treated with either DMSO or FASN inhibitor (GSK2194069; N=3 experimental replicates), with each data point representing the average of three technical replicates per experiment. Infection was normalized to the no-PF74 condition within each group, and data were fit with a 4-parameter non-linear regression with variable slope. 95% confidence intervals are shown with dashed lines centered around the mean fit for each curve. (H) Comparison of the inhibitory concentration 50% (IC50) between cells that were pretreated with FASN-inhibitor was calculated based on per-experiment 4-parameter regressions from (H). Statistical significance was determined by paired t-test. (I) Schema of mechanistic targets of experimental compounds utilized in primary CD4+ T cell experiments for (K-P). (J) The impact of FASN inhibition (GSK2194069; 25uM) on the efficacy of the antiretroviral Raltegravir in activated primary CD4+ T cells, which targets viral integration, was evaluated (N=3 donors), with each data point representing the average of three technical replicates per experiment. Infection was normalized to the no-antiretroviral condition within each group, and data were fit with a 4-parameter non-linear regression with variable slope. 95% confidence intervals are shown with dashed lines centered around the mean fit for each curve. (K) Comparison of the IC50 of Raltegravir with and without FASN inhibitor was calculated based on per-experiment 4-parameter regressions from (E). Statistical significance was determined by paired t-test. (L-M) As with K-L, but with integrase inhibitor, Efavirenz. (N-O) As with K-L, but with PF74, which targets nuclear translocation. *Statistical Significance:* *p<0.05, **p≤0.01, ***p≤0.001, ****p≤0.0001. *Abbreviations*: DMSO = dimethyl sulfoxide; FASNinh = FASN inhibitor; IC50 = inhibitory concentration 50%;

To quantify these changes to the nuclear envelope, we manually segmented the inner and outer nuclear membranes in each image, as well as a midline between them. With this approach, we were able to take repeated measurements of the perinuclear width at set distances along each segment (see *methods)*. This analysis resulted in a per-cell distribution of width measurements which could be statistically characterized. Comparison of the mean width of the perinuclear space revealed a statistically significant widening (**Figure 4D**) under FASN inhibition, which was prevented by exogenous fatty acids. Additionally, the nuclear envelope was significantly more variable during FASN inhibitor treatment compared to either the DMSO control or exogenous fatty acid addback condition (**Figure 4E**). These changes in the perinuclear width were so profound that on average ∼30% of measurements exceeded the 95^th^ percentile of width measurements in the DMSO-control (**Figure 4F**). Alterations in the barrier between the cytosol and the nucleus were seemingly independent of nuclear content as the total size of the nucleus was unchanged in these cells (**Figure S4C**).

Since previous studies established a role for mitochondrial metabolism in HIV-1 susceptibility^4,12,51^, we examined mitochondrial morphology and content during FASN inhibition as well. We did not observe differences in the mitochondria, which displayed equivalent morphology (**Figure S4D**) and total number per cell (**Figure S4E**). As mitochondrial oxidative phosphorylation depends on the structural integrity and organization of the inner mitochondrial membrane, we also examined whether FASN inhibition altered mitochondrial cristae, where respiratory supercomplexes form^52^. Yet again, we could not identify any differences between treatment groups (**Figure S4F**). Taken together, these data identify the nuclear envelope as selectively disrupted by FASN inhibition in T cells and nominate its remodeling as a potential determinant of lipid-dependent HIV-1 restriction.

HIV-1 capsid gains access to the nucleus by translocating through nuclear pores^46^. To test if alterations in the morphology of the nuclear envelope permitted pore-free capsid entry, we pre-treated CEM-GXR cells with PF74, which targets the nuclear translocation of HIV-1 capsid through nuclear pores^53–56^. PF-74 successfully blocked infection under both DMSO-treated and FASN inhibited CEM cells, indicating that translocation was occurring through nuclear pores. Strikingly, PF74 was ten-fold more effective at restricting infection in the presence of the FASN inhibitor GSK2194069 compared to PF74 alone (**Figure 4G-H**). These data suggest that FASN inhibition alters how HIV-1 interacts with the nuclear pore, raising infection at baseline but potentiating the activity of anti-retroviral drugs that target this step.

We next tested how selective the effect of concurrent FASN inhibition on PF74 was in primary activated CD4+ T cells. FASN inhibition had no effect on the efficacy of antiviral compounds that target other stages of the viral lifecycle (schema of antiretrovirals in **Figure 4I**). As shown in **Figure 4J-K**, Raltegravir^15,16^, an inhibitor of HIV-1 integration, inhibited infection in a dose-dependent manner but had no interaction with FASN inhibition, exhibiting no significant difference in IC50 with or without FASN inhibition across primary donors. The same was true for Efavirenz^57^ (**Figure 4L-M**), which targets reverse transcription, and for Darunavir^58^, an inhibitor of particle maturation which impacts the viral lifecycle after the fluorescent readout of infection assay (**Figure S4G**). As with CEM-GXR cells, PF74 potency was enhanced by FASN inhibition in primary CD4+ T-cells (**Figure 4N-O**). These data support a model by which FASN-dependent lipid synthesis safeguards the integrity of the nuclear envelope as a barrier to capsid translocation through nuclear pores and reveal that metabolic factors can be harnessed to potentiate antiviral therapies.

## DISCUSSION

In this work, we reveal that the capacity to synthesize fatty acids restricts HIV-1 infection by regulating normal architecture of the nuclear envelope. From a metabolically focused small molecule screen, inhibitors targeting key enzymes that catalyze stepwise *de novo* synthesis of fatty acids significantly increased the proportion of CD4+ T cells with integrated virus. These enzymes synthesize palmitate, which serves as the precursor for all endogenously produced fatty acids in CD4+ T cells. In the absence of FASN activity, CD4+ T cells remodel the phospholipid pool to contain longer, unsaturated fatty acids that disrupt the structure of the nuclear envelope. The nuclear envelope is a major checkpoint in the HIV-1 lifecycle, as the intact capsid must pass through the nuclear pore for infection to occur ^reviewed in^ ^59^. While we expect that FASN may similarly restrict other viral pathogens with nuclear lifecycle stages, such as Epstein-Barr virus and other herpesviruses, fatty acid synthesis may play alternative roles for pathogens that primarily engage other cellular architectures that are constructed from fatty acids. As examples, dengue virus remodels the membrane of the endoplasmic reticulum to support new virion assembly, while rotaviruses instead hijack lipid droplets to generate viral replication factories^60–62^. This study therefore highlights how host cell metabolism can sustain the intracellular structures which must be transversed or utilized by viral pathogens to productively infect host cells and establish viral disease.

We also report a comprehensive, high-throughput screen for metabolic pathways that regulate cellular susceptibility to a chronic viral infection. Previous studies have shown that HIV-1 preferentially infects CD4+ T cells with a greater capacity for mitochondrial fuel oxidation^4,11,12^, which led us to expect that the screen would predominantly yield repressors of infection. Instead, 25 of the 27 active compounds enhanced infection. This unexpected bias suggests that metabolic stress can create a state of heightened susceptibility, even though the virus still relies on the host cell to meet its bioenergetic and biosynthetic demands. Several stress pathways were identified that, to our knowledge, have not been previously associated with infection susceptibility, including inhibitors that drive ER stress as well as those that prevent cells from regenerating reduced glutathione and NADPH for detoxifying reactive oxygen species. Whether this permissive state is caused by cellular damage or by the adaptive response to restore homeostasis remains an open question for future investigation.

An important distinction is that FASN-dependent restriction operates differently from other canonical anti-viral restriction factors. Proteins such as APOBEC3 family members and MX2 interrupt defined viral processes^5–10^, whereas fatty acid synthesis restricts infection by sustaining the physical organization of the nuclear envelope, a constitutive cellular structure through which the capsid must pass. The strength of this barrier is therefore environmentally responsive, as cells can bypass the requirement for *de novo* synthesis when suitable extracellular fatty acids are available, namely saturated long-chain fatty acids such as palmitate or stearate. Susceptibility is therefore not entirely cell-intrinsic, instead emerging from the interaction between the T cell and the composition of lipids within its local environment. This means that variation in lipid availability across tissue microenvironments and individuals has the potential to influence the efficiency of HIV-1 infection^63–66^. More broadly, the consequences of metabolic stress for infection may depend on the resources available to the host cell in the local environment.

This study also reveals that metabolic features of host cells and their environments can be leveraged to sensitize infected cells to antiviral therapies. The finding that the potency of PF74 is increased in FASN-inhibited cells suggests that lipid metabolism can modulate the sensitivity to capsid-binding inhibitors. This is particularly important to understand clinically given the recent advent of semi-annual injections of Lenacapavir for HIV prophylaxis, as Lenacapavir binds to an overlapping site on capsid as PF74^67–69^. As Lenacapavir establishes more extensive interactions with adjacent capsid subunits compared to PF74, the capacity to further improve the efficacy of this antiviral with FASN inhibition will need to be carefully tested. FASN-inhibitors are currently being clinically investigated for the treatment of metabolic dysfunction-associated steatohepatitis (MASH)^70^. Given the high prevalence of MASH among people living with HIV^71^, it will be important to determine whether the potentiation of capsid-targeting antiretrovirals exceeds the impact of FASN inhibition on cellular susceptibility to HIV-1 *in vivo*, both in the context of treatment and prophylaxis.

In sum, the data presented in this study position the cellular metabolic environment as an additional layer of the virus-host interface that works in conjunction with genetically encoded restriction and susceptibility factors. We demonstrate that metabolic features of host cells have the potential to impact the intracellular structures and architectures that viral pathogens must navigate, and how identifying these interfaces can reveal unexpected ways to potentiate antiviral therapies.

### Limitations

In this study, we primarily utilized chemical inhibitors to perturb lipid metabolism in T cells. While an off-target impact of these inhibitors is possible, we observe consistent infection phenotypes upon targeting multiple steps of fatty acid synthesis, as well as for multiple inhibitors against FASN. That this HIV-1 susceptible state can be prevented by the addition of the end-product of fatty acid synthesis provides further evidence that this pathway restricts HIV-1 infection. We also focus on how FASN and the capacity for lipid synthesis regulate early events in the viral lifecycle through genome integration. However, productively infected cells *in vivo* also produce new virions, where FASN inhibition has been previously shown to reduce particle production, likely reflecting the need to acquire a lipid envelope from the host cell^72^. Thus, the net effect of FASN inhibition during multicycle infections remains unresolved. However, even if particle production is impaired by FASN-inhibitors *in vivo*, it is the earliest infection events that seed the persistent viral reservoir. Therefore, understanding factors that impact the early steps of the viral lifecycle may be just as important, if not more so, as those that impact viral spread.

## METHODS

### Cell maintenance and culture

CEM-GXR cells were maintained in RPMI1640 (Gibco, #11875093) supplemented with 10% heat inactivated FBS (R&D Systems, #S11550) 100 units/mL penicillin/streptomycin (Gibco, #15140122) and 25mM HEPES (Gibco, #15630080). Activated primary human CD4+ T cells were maintained in the same media, supplemented with 10 ng/ml IL-2 (PeproTech, #200-02-50UG). HEK293T cells (ATCC, #CRL-3216) utilized to propagate viral stocks were procured from ATCC and expanded once prior to cell freezing for transfection grade cell stocks in standard DMEM (Gibco, #11995073) supplemented with 10% heat inactivated FBS and 100 units/mL penicillin/streptomycin. All cells were cultured at 37°C in a humidified incubator containing 5% CO_2_.

### Generation of plasmid stocks for viral transduction

HIV-1 NL4-3 ΔEnv GFP Reporter Vector (NIH-AIDS Reagent program, #11100), VSVG expression vector (pMD2.G Addgene #12259), and HIV^89.6^ expression vector (NIH-AIDS reagent program, #ARP-3552) were first spread on agar plates supplemented with 100ug/ml Carbenicillin (Thermo Fisher Scientific, #455360050) to isolate single colonies, which were then expanded into liquid cultures for isolation by ZymoPure II plasmid maxiprep (Zymo Research, #D4202). Plasmids were grown at 30°C, with DNA concentration and purity determined on a NanoDrop microvolume spectrophotometer. All plasmids were sequence verified by full length plasmid sequencing with Plasmidsaurus.

### Generation of viral stocks

For the generation of viral stocks, HEK293T cells were seeded in standard DMEM growth media (see cell maintenance and culture) overnight until cells were ∼70% confluent. For VSVG-NL43 ENV-GFP, and HIV^89.6^ stocks were generated via transduction of HEK293T cells with genome encoding plasmid expression platforms. For VSVG-pseudotyped viral stocks, 7.5 µg of VSVG-NL43 ENV-GFP plasmid and 64.5 µg of VSVG expression plasmid were co-transduced. For HIV^89.6^, 7.5 µg of HIV^89.6^ plasmid was transduced in a BL2+ research setting. DNA was first mixed with 3 mL of OPTIMEM reduced serum media (Gibco, #31985062) prior to being supplemented with Polyethyleneimine (LifeTechnologies, #BMS1003) at a 3:1 volume to DNA mass ratio. After a 15-minute pre-incubation, PEI: DNA complexes were applied dropwise to HEK293T cell monolayers. Transduction mixtures were removed after 3 hours of incubation at 37°C and replaced with pre-warmed DMEM growth media (without antibiotics). After 48 hours, cell supernatants were collected by centrifugation and were then further purified with a 45 µm syringe filter. Viral stocks were then aliquoted at stored at -80°C until further use.

### Titration of viral stocks

For VSVG-NL43 ENV-GFP, virus was tested by dilution assay on CEM-GXR cells in a format mimicking infection experiments. 50,000 CEM-GXR cells were cultured in 96-well plates for 24 hours to simulate the addition of a metabolic inhibitor. After this 24-hour preincubation, viral stocks were titrated on CEM-GXR cells, and infection was quantified 72 hours later by flow cytometric detection of GFP expression among live cells. A viral dilution where ∼10% infection was achieved in CEM cells was determined and utilized for following experimentation.

For HIV89.6, viral titers were quantified using the TZM-bl assay^73,74^. TZM-bl cells were seeded at 300 K/mL density in complete DMEM (Dulbecco’s Modified Eagle Medium supplemented with 10% fetal bovine serum, 1% penicillin/streptomycin, and 25 mM HEPES) containing 75 µg/mL DEAE-dextran (Sigma Aldrich, #D9885-10G) in a flat clear bottom plate (Corning, #3904). Cells then infected by serially diluted HIV^89.6^ viral stocks at 2x concentration. This generated a final cell density of 150K/mL, and titration started at 1:50 dilution and 3-fold dilution between each concentration. Cells were spin-inoculated at 1000 rpm for 3 minutes and infection continued at 37°C and 5% CO_2_ for 48 hours. For luciferase reaction, cells were pelleted and resuspended in 1x OLST buffer (20 mM Tris, 1.07 mM MgCl_2_, 2.67 mM MgSO_4_, 100 µM EDTA, 250 µM ATP, 17 mM DTT, 1% (v/v) Triton-X, and 25mM D-luciferin). Luciferase activities were measured by a PHERAStar microplate reader immediately after a final spin at 1500 rpm for 3 minutes once 1x OLST buffer was added.

### Small molecule inhibitors

All compounds were reconstituted in DMSO at a stock concentration of 10 mM for downstream use. The small molecules utilized in this study outside the context of the small-molecule screen were as follows: ND630 (Firsocostat; MedChem Express #HY-16901), GSK2194069 (MedChem Express #HY-12325), TVB2640 (Denifanstat; MedChem Express #HY-112829), PF-04620110 (MedChem Express #HY-13009), A-922500 (MedChem Express #HY-10038), PF74 (PF-3450074; Medchem express #HY-120072), Raltegravir (MedChem Express #HY-10353), Efavirenz (MedChem Express # HY-10572), Darunavir (MedChem Express #HY-17040).

### Small molecule screen

#### Human primary CD4+ T cell *ex vivo* activation, expansion, and freezing

Frozen primary human CD4+ T cells from two healthy donors were purchased (StemCell Technologies, #70026, Donor IDs CE000645 and CD0010557). Cells were rested overnight at 1×10^6^ cells/mL density in complete RPMI 1640 (RPMI 1640 supplemented with 10% fetal bovine serum, 1% penicillin-streptomycin, and 25 mM HEPES). The next day, cells were stimulated using Dynabeads (Thermo Fisher Scientific, #11161D) at a 1:1 ratio in complete RPMI supplemented with 10 ng/mL human recombinant IL-2 (PeproTech, #200-02-50UG) for three days. Activated cells were removed from the Dynabeads and re-cultured in RPMI for expansion. The density of cells was monitored daily. For 8 additional days, RPMI supplemented with IL-2 were added to maintain a 0.75×10^6^/mL density. Expanded CD4+ T cells were frozen in CryoStor CS10 freezing media (StemCell Technologies, #07930) at 60×10^6^/mL density according to the manufacturer’s instructions.

#### High-throughput screening

Ludwig metabolic library 2 was provided by ICCB-Longwood screening facility^75^. This library contains 10 plates, with 24 compounds each plate and 10 concentrations (ranged from 10 mM to 500 nM) per compound, generating 10-point titration curves for all 240 compounds. Compounds were purchased from Sigma, Selleck Chemicals, Toris, and Cayman Chemicals, and reconstituted in dimethyl sulfoxide (DMSO). Frozen expanded primary CD4+ T cells generated as described above were thawed and rested overnight at 1×10^6^ /mL density in complete RPMI. The next day, cells are first stained with CellTrace Violet proliferation dye (Invitrogen, #C34557) according to manufacturer’s instruction and then stimulated using ImmunoCult human CD3/28 T cell activator (StemCell, #10971) at a concentration of 25 µL per 1×10^6^ cells in complete RPMI supplemented with IL-2 for 24 hours at 37°C. After stimulation, cells were pelleted and resuspended at 1×10^6^ cells/mL density in fresh complete RPMI supplemented with IL-2 to prevent further stimulation. For pre-treatment with inhibitors, compounds from the source library plates were transferred to polystyrene U-bottom 384-well plates (Greiner BIO-ONE, #787979) via a Beckman Coulter Echo 655 acoustic liquid dispenser. Cell suspension was transferred to the 384-well plates containing compounds via a Thermo Multidrop Combi rapid plate dispenser to generate a final concentration range of 33 µM to 1.67 nM for each compound. After 24 hours of inhibitor treatment at 37°C, cells were infected with NL43ΔENV GFP-VSVG viruses and spin-inoculated at 300x g for 5 minutes. Cells were infected for 72 hours before harvesting. Cells were washed twice with sterile PBS. Staining for surface markers staining and live/dead discrimination were performed concurrently in FACS buffer (1% BSA, 1 mM EDTA in 1x PBS) at 1:200 dilution for surface markers and 1:1000 for the live/dead dye on ice in the dark for 30 minutes. The following antibodies were used for surface marker and live/dead staining: Brilliant Violet 650 mouse anti-human CD4 antibody (BD Biosciences, #740563), PE-Cyanine 5 mouse anti-human CD62L (Invitrogen, #50-140-71), LIVE/DEAD Fixable Near-IR stain (Thermo Fisher Scientific, #L10119). Cells were fixed in 2% PFA (Thermo Fisher Scientific, #J19943.K2), which was diluted from 4% to 2% in FACS buffer and analyzed using the Intellicyt iQue Screener PLUS.

#### CD4+ T cell immunophenotyping

For memory subsets and helper T cell subsets immunophenotyping, unstimulated and unexpanded CD4+ T cells were first stained with LIVE/DEAD Fixable Near-IR stain (ThermoFisher Scientific, 525 L10119) diluted in 1X PBS (ThermoFisher, 14190-250) according to manufacturer’s instructions. Chemokine receptors (CXCR3, CCR6, and CXCR5) staining were performed in 2% FBS diluted in PBS at a 1:20 dilution for CXCR3 and CXCR5, and a 1:40 dilution for CCR6 at 37°C for 20 minutes. Extracellular staining was performed at RT for 20 minutes. Intracellular staining was performed using eBioscience FOXP3 and transcription factor staining buffer set (ThermoFisher Scientific, #00-5523-00) according to manufacturer’s instructions. Antibodies for extracellular markers were used at 1:200 dilution (except CD25 were stained at 1:50 dilution), while antibodies for intracellular targets were used at 1:100 dilution. Cells were analyzed using a BD FACSymphony A5 cytometer.

The following antibodies were used for immunotyping CD4+ T cells via flow cytometry: RealBlue 705 mouse anti-human CD3 antibody (BD Biosciences, #757064), Brilliant Ultra Violet 395 mouse anti-human CD4 antibody (BD Biosciences, #564724), Brilliant Violet 711 mouse anti-human CD45RA antibody (BioLegend, #304138), Brilliant Ultra Violet 737 mouse anti-human CD62L antibody (BD Biosciences, #568329), Brilliant Violet 421 mouse anti-human CXCR3 antibody (BD Biosciences, #562558), Brilliant Violet 510 mouse anti-human CCR6 antibody (BD Biosciences, #563241), Alexa Fluor 647 mouse anti-human FOXP3 antibody (BioLegend, #320114), Alexa Fluor 488 rat anti-human CXCR5 antibody (BD Biosciences, #558112), and PE mouse anti-human CD25 antibody (BioLegend, #BC96).

### Infection screening in CEM-GXR and Primary CD4+ T cells

50,000 CEM-GXR cells or 24-hour activated primary CD4+ T cells were plated in 96-well plates in a total volume of 100 μl. For primary cell experiments, recombinant human IL-2 was supplemented to media at 20 ng/ml concentration. Metabolic inhibitors were prepared at a 2x final concentration in the same media conditions and added 1:1 to cells to achieve a final 1x concentration of both metabolic inhibitors and IL-2. After preincubation overnight, cells were infected with NL43ΔENV GFP-VSVG at concentrations described in the method detail for *viral titration*. Primary cells were inoculated via centrifugation at 350x g for 10 minutes to facilitate infection. After 72 hours, cells were harvested by centrifugation (350x g for 3 minutes at RT), and washed 1x with sterile PBS. Cells were then stained with a 1:1000 dilution of LIVE/DEAD Fixable Near-IR stain (ThermoFisher Scientific, 525 L10119) for 10 minutes at room temperature in the dark. Cells were then either fixed 2% PFA (Thermo Fisher Scientific, #J19943.K2, diluted from 4% to 2% in FACS buffer) or stained for surface expression of CD4 using a 1:200 dilution of Brilliant Ultraviolet 395 mouse anti-human CD4 antibody in FACS buffer (BD Biosciences, #564724) prior to fixation as above. For primary cell experiments with wildtype virus, cells were fixed for 1 hour utilizing the FOXP3 cell fixation kit (Invitrogen, #00-5523-00), permeabilized for 15 minutes at room temperature, and stained for p24 expression (Beckman Coulter, Clone KC57# 6604667) at a 1:200 dilution for 1 hour at room temperature. After washing cells 3x in permeabilization buffer, cells were then reconstituted in PBS for collection by flow cytometry.

For CEM-GXR experiments, viral GFP was utilized to detect infection in live cells. For primary human experiments with pseudovirus, either GFP was utilized alone, or in conjunction with p24 expression as indicated in text. For wildtype infections, the concurrent downregulation of CD4 and expression of p24 was utilized to define infected cells. All flow cytometry was collected on either a BD FACSymphony or LSRFortessa, and data analysis was conducted in FlowJo (Version 10.10.0).

### Stable isotope tracing of ^13^C_6_-glucose into palmitate

#### Cell culture and metabolite extraction

CEM-GXR cells were plated in 6-well plates in technical triplicate at a seeding density of 1×10^6^ cells/ mL in RPMI1640 that lacked glucose (Gibco, #11879020). Cells were first treated with 25 µM of ND630, GSK2194069 or a volume matched DMSO control. ^12^C-glucose (Thermo Fisher Scientific, #A2494001) or ^13^C_6_-glucose (Cambridge Isotope Laboratories, # CLM-1396-2) was supplemented at matched concentration to that in normal growth media (11.1 mM) and incubated at 37°C overnight. One plate at a time, cells were removed from the incubator and 1.5 mL of cells per condition were washed in cold 0.9% (w/v) HPLC-grade NaCl (Thermo Fisher Scientific, #446212500) in OMNISolv LC-MS grade water (VWR, #EM-WX0001-1). Cells were then resuspended in 300 µl of ice-cold LCMS-grade methanol (MeOH; VWR, #BDH85800.400) containing 0.02% (w/v) Butylated hydroxytoluene (BHT) (Sigma, #PHR1117-1G) and 1.33 µg/mL of Cis-Heptadeanoic acid internal standard (Caymen Chemical, #36254) to extract metabolites. 1 mL of ice-cold Methyl tert-butyl ether (MTBE) (SigmaAldrich, #34875) was added, and cells were then mixed end-over-end for 15 minutes at 4°C prior to the further addition of 250 µl/sample of ice-cold HPLC grade water. Samples were again mixed end-over-end for 10 minutes at 4°C and centrifuged at 3000x g for 5 minutes at 4°C to induce phase separation. The non-polar fraction (top-layer) was harvested and placed into amber glass tubes for derivatization of fatty acids into fatty acid methyl esters (FAMES).

#### Derivatization of fatty acids into FAMES

Non-polar fractions were dried under nitrogen gas on a 48-position MULTIVAP nitrogen evaporator (Organomation, #11848) and then dissolved in 100 µl toluene (Sigma Aldrich, #650579-1L). 200 µl of 2% sulfuric acid (Sigma Aldritch, #1120800510) in HPLC-grade methanol was then added. Samples were vortexed and incubated overnight at 50°C. Samples were then supplemented with 500 µl /sample of 5% NaCl (w/v) (Thermo Fisher Scientific, #446212500) in LC-MS grade water. 500 µl/sample of HPLC-grade hexane was added (Fisher Scientific, #AA39199K2) and the non-polar layer was isolated. This last extraction was repeated, and the hexane isolates were merged in amber glass tubes per sample and dried under nitrogen gas. Dried FAMES were finally reconstituted in 50 µl hexane per sample and transferred to glass autosampler vials for GC-MS analysis.

#### Sample collection by GC-MS

Gas chromatography/Mass spectrometry (GC/MS) was conducted on an Agilent 5977C GC/MSD (Agilent, G7077C) fit with an DB-Fast FAMES column under Helium gas carrier flow (1.764 mL/min,). 1 µl of FAMES were injected into the inlet (250°C) by a Gerstel MultiPurpose Robotic sampler (#RSI717388). The GC oven was set to 50°C for 0.5 minutes and then ramped at 25°C/min until 194°C at which it was held for 1 minute. The oven was then ramped at a rate of 5°C/min. until achieving 245°C, where it was held for a final 3 minutes. The MS source/quadrupoles were held at 230°C and 150°C respectively. Following a 3-minute solvent delay, the instrument scanning range was set between 70 and 412 m/z, with a scan frequency of 4.1 scans/sec and a 0.1m/z step size. 1 µl of hexane was utilized to wash the column between every three samples.

### Isolation of CD4+ T cells from healthy donors

Human CD4+ T cells from healthy blood donors were collected from the Massachusetts General Hospital Blood Transfusion Service under IRB protocol #2005P001218, which permits the use of blood for research purposes. Buffy coats were first diluted 1:1 with sterile PBS, and PBMCs were subsequently isolated via density centrifugation over Lymphoprep (StemCell Technologies, #07811) in SepMate PBMC isolation tubes (StemCell Technologies, #85450). CD4+ T cells were then isolated from PBMCs through negative selection, using the EasySep Human CD4 negative selection kit (StemCell Technologies, #17952) according to manufacturer’s protocols. CD4+ T cells were then either stimulated overnight with Immunocult CD3/CD28 activator solution (Metabolic drug screen only; StemCell Technologies, #10991) or with washed, anti-CD3/CD28 dynabeads at a 1:1 bead to cell ratio (Gibco # 11161D) in the presence of 10 ng/ml recombinant human IL-2. After 24 hours of activation, activation signal was removed. For Immunocult activator, cells were washed 1x with PBS. For anti-CD3/CD28 dynabeads, activator beads were removed by magnetic separation. Isolated cells were then utilized in downstream assays or expanded in RPMI1640 supplemented with 10% heat inactivated FBS, 100 units/mL penicillin/streptomycin and 25 mM HEPES in the presence of 10 ng/mL recombinant human IL-2 for up to 8 days prior to freezing for future re-stimulation.

### Conjugation of fatty acids with BSA

A 1.36 mM stock of Fraction V fatty acid free BSA (Gold Bio, #A-421-100) in 150 mM NaCl was prepared in a glass beaker in a 37°C water bath. Fatty acid stocks were reconstituted in 150 mM NaCl and heated to 70-90°C in a water bath and mixed until dissolved. BSA-NaCl was added to dissolved fatty acids at a 1:6 molar (BSA: FA) ratio and incubated at 37°C for 1 hour to facilitate conjugation of fatty acids to BSA. For fatty-acid free BSA controls, this same process was repeated but BSA-NaCl was added to NaCl at matched volumes as fatty acids. Fatty acid stocks were then pH balanced to 7.4 utilizing 1N NaOH, sterile filtered and then stored at -20C in glass tubes until further use. Fatty acids utilized in this work are listed, with the stock concentration indicated per fatty acid: Sodium palmitate (C16:0; 4 mM; Sigma P9767-5G), Palmitoleic acid (C16:1; 4mM; Fisher scientific AC376912500), Sodium Stearate (C18:0; 2 mM; S3381-5G), Sodium Oleate 97.0+%, TCI America™ (C18:1; 4 mM; FisherScientific O005725G), Linoleic acid sodium salt (C18:2; 4 mM; Sigma L8134-500MG), dihomo-gamma-linolenic acid (C20:3; 1 mM; Abcam ab145213) were conjugated for this work at the indicated working concentrations. For assays utilizing these fatty acids, all stocks were first diluted to 0.8 mM in growth media to prevent changes in BSA concentration from impacting downstream analysis.

### Palmitate uptake assays with click chemistry

Fatty acid uptake was evaluated based on the uptake of palmitic acid-alkyne (Cayman Chemical, #13266), which is compatible with click chemistry^34–36^. Briefly, 50,000 acutely activated primary human T cells were plated in two 96-well plates in 200 µl culture volume and pre-treated with either DMSO, FASNinh (GSK2194069 at 50 µM) or FASNinh with exogenous palmitate-BSA/oleate-BSA (50 µM) and incubated overnight. Following, cells were washed 1x in PBS and then stained with 1:1000 fixable near-IR dead cell stain kit (Invitrogen, #L10119) for 10 minutes at RT. Stain was washed off once with HBSS (Thermo Fisher Scientific, # 14025092) and cells were then resuspended in 50 µl /well of HBSS. 96-well plates were incubated for 15 minutes at either 37°C or 4°C to facilitate or inhibit uptake respectively. In parallel, palmitate-alkyne (50 mM stock in DMSO) or palmitate-BSA (4 mM stock in NaCl; see *method* for conjugation of fatty acids to BSA) were diluted in HBSS to a concentration of 100 µM (2x final concentration) and were prewarmed or chilled to match cell culture conditions. 2x fatty acid solutions were quickly added to the appropriate cell condition and incubated at their respective temperatures, generating the following: 37°C + palmitate-alkyne, 37°C + palmitate-BSA, 4°C + palmitate-alkyne. Cells were co-incubated with fatty acids for 30 minutes, after which cells were immediately fixed with eBiosciences FOXP3/transcription factor kit (Invitrogen, #00-5523-00) for 30 minutes. Cells were then washed twice in 1x permeabilization buffer from the kit, diluted in diH2O. Cells were then permeabilized for 20 minutes in permeabilization buffer, after which they were resuspended in click-mix containing 1 mM copper sulfate (CuSO_4_; Honeywell, #C1297-100G), 10 mM sodium ascorbate (Sigma-Aldrich, #A7631), 10 mM tris-hydroxypropyltriazolylmethylamine (THPTA; Click Chemistry Tools, #1010-100), 10 mM Aminoguanidine (Caymen Chemicals, #81530), and 5 µM AZDye647 Azide (Vector Laboratories, #CCT-1299-5) for 1 hour at room temperature. Cells were washed 1x with FACS buffer, incubated in FACS buffer for 30 additional minutes to reduce non-specific labeling of cells, and then resuspended in PBS for analysis by flow cytometry.

### Lipidomics

Lipidomic analysis of CEM-GXR cells was conducted in collaboration with the UCLA Quantitative Lipidomics core facility. Approximately 1×10^6^ cells per sample were transferred to extraction tubes with PBS. A modified Bligh and Dyer extraction (Hsieh, 2020) was carried out on all samples. Prior to biphasic extraction, a standard mixture of 74 lipid standards was added consisting of Avanti Ultimate Splash ONE mix (Avanti, #330820) with Splash Booster (Avanti, #330740). Following two successive extractions, pooled organic layers were dried down in a Thermo SpeedVac SPD300DDA using ramp setting 4 at 35°C for 45 minutes with a total run time of 90 minutes. Lipid samples were resuspended in 1:1 methanol/dichloromethane with 10 mM Ammonium Acetate and transferred to robovials (Thermo Fisher Scientific, #10800107) for analysis. Samples were analyzed on the Sciex 5500 with DMS device and an expanded targeted acquisition list consisting of 1450 lipid species across 17 subclasses. Differential Mobility Device was tuned with EquiSPLASH LIPIDOMIX (Avanti, #330731). Data analysis performed on an in-house data analysis platform comparable to the Lipidyzer Workflow Manager^76^. Instrument method including settings, tuning protocol, and MRM list available in Su et. al. Quantitative values were normalized to cell number. Lipid species with odd-chain carbon patterns that are not relevant to *in vitro* mammalian cell culture were filtered prior to data analysis (see statistical analyses section for more details).

### Transmission electron microscopy (TEM)

TEM was conducted at the Harvard Medical School electron microscopy core facility. Unless noted otherwise, all reagents were procured from electron microscopy sciences. CEM-GXR cells were pelleted by centrifugation and fixed in a solution of 2.5% Glutaraldehyde 1.25% Paraformaldehyde (Electron Microscopy Science, #15949) and 0.03% picric acid in 0.1 M sodium cacodylate (Electron Microscopy Science, #12300) buffer overnight at 4°C. Fixed cell pellets were then washed in 0.1M cacodylate buffer and postfixed with 1% Osmium tetroxide (OsO4)/1.5% Potassium ferrocyanide (KFeCN6) (Electron Microscopy Science, #19150) for one hour. Samples were then washed twice in water, once in Maleate buffer (MB; Electron Microscopoy Science, #18150), and were then incubated in 1% uranyl acetate (Electron Microscopy Science, #22400) in MB for one hour. Two washes in water were then conducted, and samples were then dried through subsequent dehydriding washes in an ethanol gradient (10 minutes each; 50%, 70%, 90%, 2×10 minutes 100%). The samples were then placed in propylene oxide for one hour and infiltrated overnight in a 1:1 mixture of propylene oxide (Electron Microscopy Science, #20412) and TAAB (TAAB Laboratories Equipment Ltd, #T022). The following day the samples were embedded in TAAB Epon and polymerized at 60°C for 48 hours. Ultrathin sections (about 60 nm) were then cut on a Reichert Ultracut-S microtome, picked up on to copper grids, stained with 0.2% lead citrate and examined in a TecnaiG² Spirit BioTWIN transmission electron microscope. Images were recorded with an AMT Nanosprint 43-MKII camera.

### Cell trace violet labeling of CD4+ T cells

Isolated primary CD4+ T cells were washed once with PBS to remove residual media components and were then resuspended at a concentration of 1×10^8^ cells/ mL in fresh PBS. CellTrace Violet (Invitrogen, #C34557) was resuspended in 20 µl of DMSO and added to cells at a 1:1000 dilution as per manufacturer’s protocols. Cells were incubated at 37°C for 20 minutes and were then washed with complete RPMI growth medium. Cells were then counted, and utilized for downstream applications, such as activation and metabolic inhibitor treatment.

### Statistical analysis

#### Data filtering and hit identifications of small molecule screen

All quality-control filtering and hit identifications of screen data were performed in R (v.4.3.3) within R studio (v.2024.12.0.467) using dplyr (v.1.1.4), tidyr (v.1.3.1), ggplot2 (v.3.5.1), forcats (1.0.0), scales (1.3.0), and purrr (1.0.2). Flow cytometry data were exported as CSV files from FlowJo v.10.10.0 and imported into R studio. Raw viability percentage were normalized to DMSO-treated controls. Raw infection percentage were transformed to normalized infection scores using the following formula: Briefly, wells with a normalized viability smaller than 90 were removed. Doses with at least a 10-fold differences in normalized infection between two donors were considered discordant and removed. Compounds with less than three doses remained after the viability and concordance filtering were excluded from further analysis. Mean normalized infection using the following formula:

Hit identification was done via applying a modified Tukey’s fence (k = 1) on mean normalized infection. Compounds with mean normalized infection above the upper fence were called enhancers, below the lower fence, repressors. Enhancers and repressors were then passed through a minimum normalized infection filter, removing compounds that only induced change in infection in one donor but failed to elicit any response from the other donor. Minimum normalized infection threshold was defined individually for each donor, as the 93.3^rd^ of all normalized infection scores of all viable and concordant doses of all compounds (corresponding to 1.5 standard deviations from the mean in a normal distribution). If no point on a dose-response curve of any donor of a compound met this threshold, this compound was removed.

### Analysis of ^13^C-Glucose incorporation into palmitate

Raw data analysis files were converted to MZML format by msConvert^77^ and were subsequently analyzed in EL-MAVIN (v0.12.1-beta). M/Z 270 was extracted from total ion chromatograms (TICs) at the retention time indicated by the Supelco 37 FAME mix (Supelco, #CRM47885) collected under the same method. Peaks were then integrated, and peak area was recorded. This process was repeated for each predicted isotopologue of palmitate-methyl ester at the same retention time. Integrated peak areas were then center scaled based on the distance from the average cell count, and natural isotope abundance correction was performed with IsoCor as previously described^78^. Finally, the percentage of total palmitate-methyl ester carbon contributed by each isotopologue was calculated and represented as fractional enrichment.

### Lipidomics and Lipid ontology analysis with LION

Lipid abundance data (normalized to cell number) was first filtered to only include even-chain fatty acids to focus the analysis on lipid species relevant to de novo synthesis (which utilizes 2-carbon additions at a time). Undetected analytes were imputed based on half-minimum replacement, while species undetected in all samples were filtered out of downstream analyses. Lipid ontology analysis was conducted on the LION/Web analysis platform^40^ based on lipid abundance for each lipid species detected in each treatment condition with the following settings per comparison: Welch’s t-test statistic was calculated from the raw abundance data between comparisons, with ranking direction set from low to high and alternative hypothesis testing set to two-tailed mode. Adjusted P-values and calculated LION enrichment values were utilized to generate downstream volcano plots for data representation.

For each phospholipid or precursor class, each analyte was summarized by the sum carbon length between the two fatty acid tails, and the sum desaturation number (number of double bonds). Abundance weighted means were calculated for each sample, whereby we calculated the sum of the abundances of each species multiplied by the total number of carbons (or double bonds) in that species. This sum was then normalized to the total abundance of the subclass (example: the sum of all measured phosphatidylcholines). To avoid distorting the denominator, non-detected lipid species were given a value of 0 for this analysis.

### Image quantification in FIJI

Nuclear envelope morphology was quantified based on scale-calibrated TEM images using a custom pipeline IN FIJI (version 2.16.0)^79^. First, nuclear segments were manually segmented into three categories: Inner membrane (facing the nucleus), Outer membrane (facing the cytosol) and a midline. The midline was sampled every 2 nm. At each measured point along the midline, a tangent was calculated based on 15 nm distances from the point along the midline and used to calculate a subsequent perpendicular line that intersected the midline. The distance between where this perpendicular line intersected the midline and the outer or inner membranes was calculated and summed to calculate the width of the envelope at that location. A distribution of width measurements was calculated per segment to generate a per-segment mean and standard deviation in width values. These per-segment values were then pooled to generate a per-cell distribution from which mean and standard deviation were calculated. A 95^th^ percentile threshold was calculated based on the pooled measurements from DMSO-controls and utilized to define the percentage of points per cell that were “expanded”, which was subsequently compared across groups. For metrics such as cell size, nuclear size, these were generated based on manual segmentation of cells or nuclei from 1400x images. For mitochondrial metrics, 4800x images were analyzed and manually counted.

### IC50 calculation and data fitting

For antiretroviral experiments, the percent infection in each sample was first normalized to the matched condition lacking anti-retroviral. For the FASN-inhibited condition, this was thus normalized to the FASN-inhibitor alone to permit relative comparison of IC50s with/without FASN-inhibitor. Data were fit utilizing the variable 4-parameter nonlinear regression functionality of GraphPad Prism (Version 11.0.1). Calculated IC50 values for each experimental replicate were utilized to compare IC50s between conditions.

## Code and resource availability

Code and critical reagents will be made available upon request.

## Data availability

Raw data from the small molecule screen and lipidomics experiment generated in this study will be made publicly available upon successful peer-review and publication of this manuscript.

## Acknowledgments

We would like to acknowledge the contributions of the following individuals and facilities:

Dr. Jennifer Smith, Dr. Patricia Szajner, Bryce Carr and the entire Harvard medical school lCCBL longwood screening facility staff for supporting the design and execution of the metabolic small molecule screen. Dr. Kevin Williams and the UCLA Quantitative lipidomics core, Dr. Maria Erickson and the Harvard Medical School Electron Microscopy core, Michael Waring and the Flow Core of the Ragon Institute of MGB, MIT and Harvard. We thank Drs. Nicolas Webb and Boris Juelg for their support on Intellicyt iQue data collection. We would also like to thank Drs. Lucy Walters and Michael Birnbaum for generously donating the VSV-G expression plasmid utilized to generate viral stocks, as well as Drs. Nicolas Galvez and Alejandro Balazs for donating the HIV^89.6^ genome plasmid.

