## Supplemental Figures for "Fatty acid synthesis restricts HIV-1 infection through regulation of the nuclear envelope in CD4+ T cells"

### SUPPLEMENTAL MATERIALS

A

| Donor | Gender | Age | Ethnicity | Smoker | Weight (kg) | Height (cm) | BMI | CMV Status | Blood Type |
|---|---|---|---|---|---|---|---|---|---|
| 1 | M | 28 | CAU | N | 86 | 180 | 26.5 | POS | O+ |
| 2 | F | 22 | CAU | N | 68 | 178 | 21.5 | NEG | O+ |

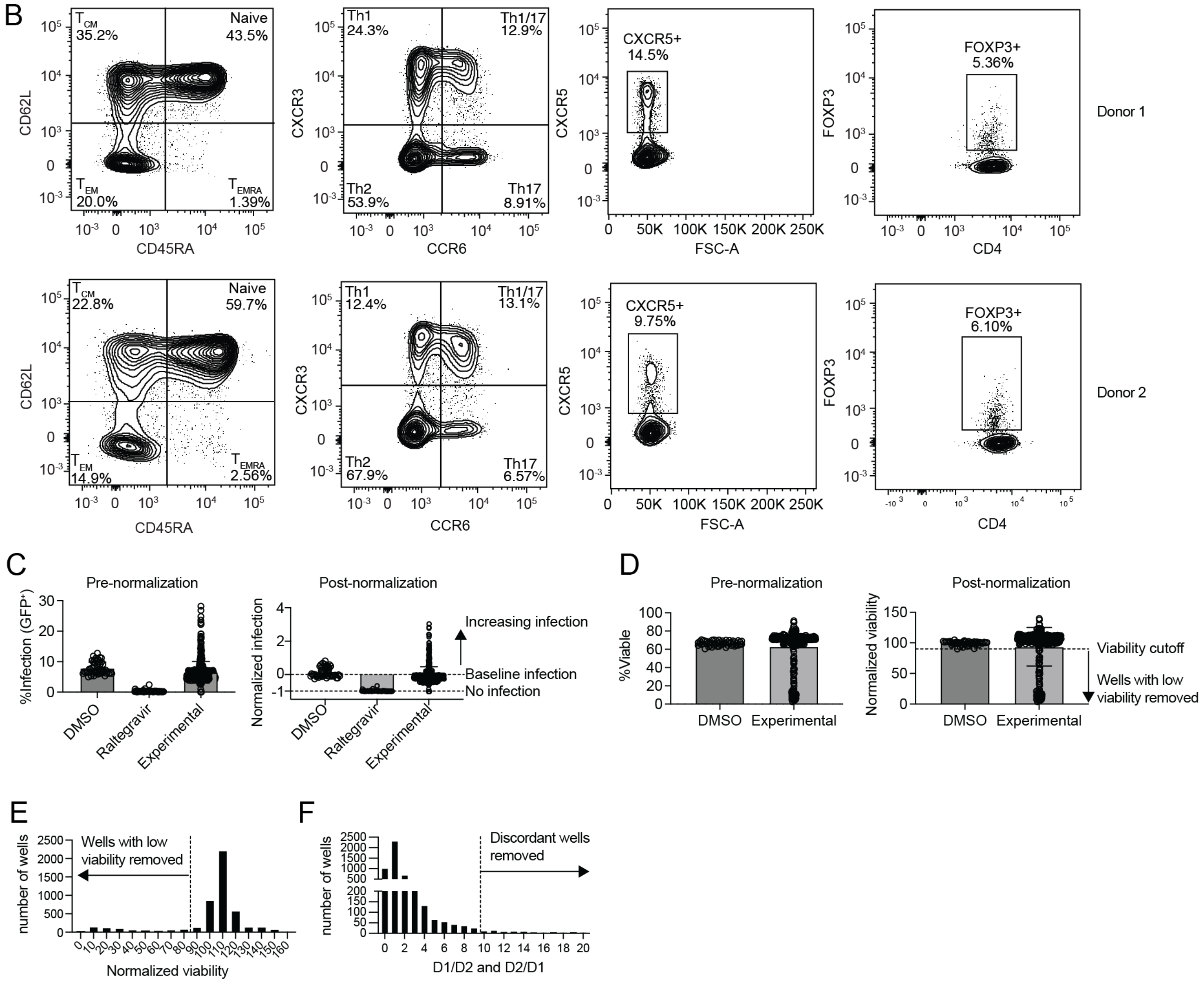


**Figure S1. (Related to Figure 1): Screen donor characterizations and data normalizations.**

(A) Summary table of donor characteristics and demographics.

(B) Immunophenotype of two human donors prior to activation and expansion. Surface markers were stained to quantify memory subsets (CD62L and CD45RA), helper T cell subsets (CXCR3 and CCR6), follicular-homing T cells ($CXCR5^+$) and regulatory T cells ($FOXP3^+$). All plots were pre-gated on single, live, CD3 and CD4 double positive T cells.

(C) Representative infection pre- and post-normalization. DMSO: DMSO-treated wells. Raltegravir: Raltegravir treated wells. Experimental: wells treated with inhibitors. Positive scores indicate higher infection over DMSO baseline. Negative scores indicate decreased infection from baseline. Bars represent mean, while error bars represent standard deviation.

(D) Representative viability pre- and post-normalization. Wells with normalized viability lower than a value of 90 (corresponding to 90% of the viability of DMSO-treated wells on the same plate) were excluded from analysis. Bars represent mean, while error bars represent standard deviation.

(E) Distribution of normalized viability of all experimental wells, highlighting most wells have normalized viability of at least 90.

(F) Distribution of ratios of normalized infection between donor #1 and donor #2. D1/D2 refers to the ratio of normalized infection of donor #1 to infection of donor #2. D2/D1 refers to the ratio of normalized infection of donor #2 to infection of donor #1.

*Statistical Significance:* $*p<0.05$, $**p\leq0.01$, $***p\leq0.001$, $****p\leq0.0001$.

*Abbreviations:* BMI = body mass index; CMV = cytomegalovirus infection history; CAU = Caucasian; POS = positive; NEG = negative; FSC-A = forward scatter area; DMSO = dimethyl sulfoxide. $T_{EM}$: T effector memory cells; $T_{CM}$: T central memory cells; $T_{EMRA}$: T effector memory re-expressing CD45RA.

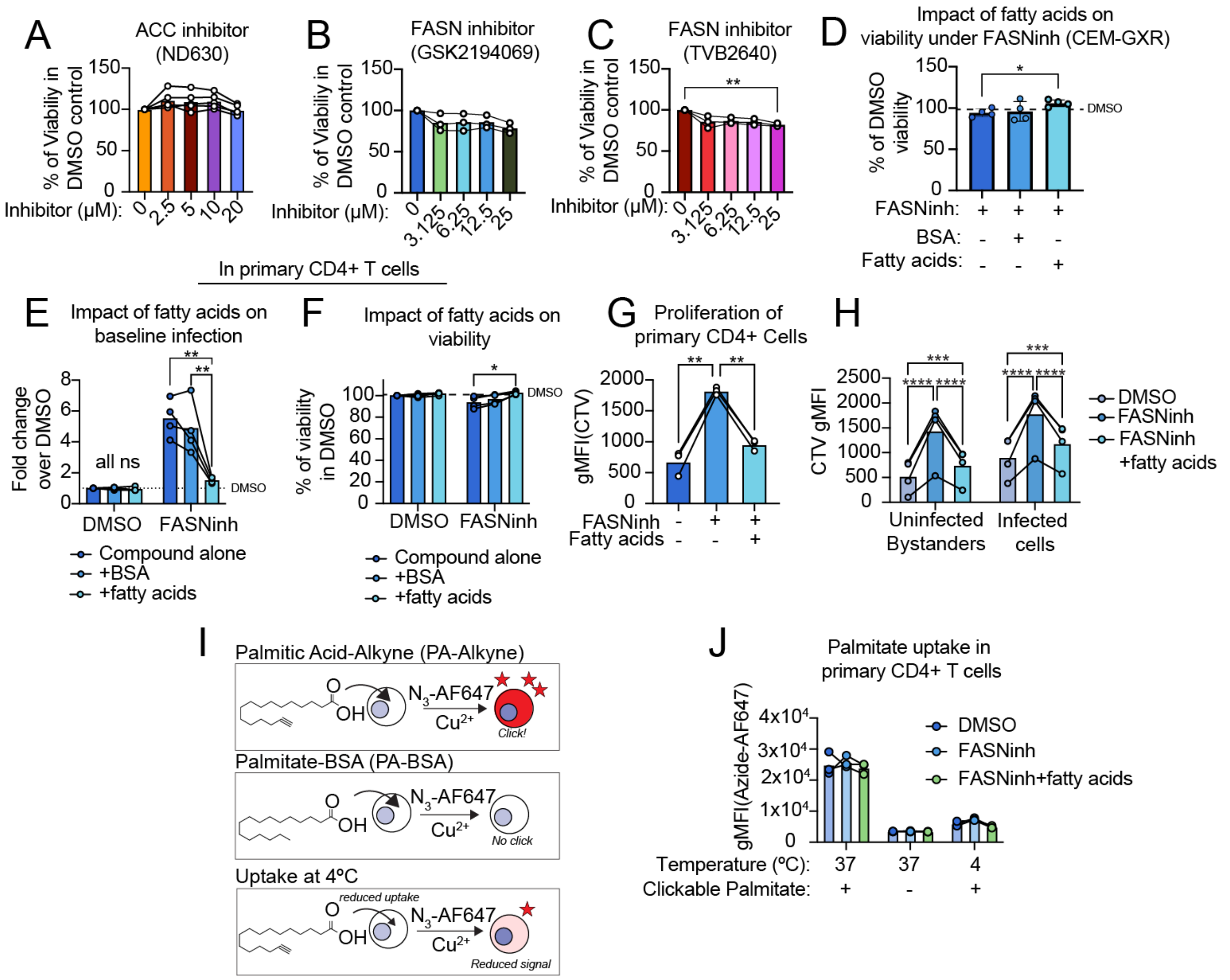


**Figure S2 (Related to Figure 2): Impact of inhibitors of fatty acid synthesis on cellular viability.**

(A) Impact of ACC inhibitor (ND630) on the viability CEM-GXR cells. Each data point represents an experimental replicate (N=4), which is the average of three technical replicates per experiment. Statistical significance was determined by two-way ANOVA with Dunnet's multiple comparison test against the DMSO-treated control.

(B-C) As with (A), but with FASN-inhibitor GSK2194069 (B; N=3 experimental replicates) or FASN-inhibitor TVB2640 (C; N=3 experimental replicates).

(D) The impact of fatty acid-free BSA or exogenous fatty acids on the viability of CEM-GXR cells under FASN inhibition (GSK2194069) was evaluated. N=4 experimental replicates, with datapoints representing the average of three technical replicates per experiment. Statistical significance was determined by two-way ANOVA with Dunnet's test for multiple comparisons against the FASN-inhibited condition. Data are represented as the percent of viability on per-plate DMSO controls.

(E-F) Evaluation of how exogenous fatty acids or fatty-acid free BSA contribute to HIV-1 infectivity (E) or cell viability (F) in either DMSO-treated or FASN-inhibited activated primary CD4+ T cells (N=3 primary donors). Statistical significance was determined by two-way ANOVA with Dunnet's test for multiple comparisons against the DMSO or FASN-inhibited control in each group.

(G) The proliferation of activated primary CD4+ T cells pretreated was assessed based on the retention of Cell Trace Violet across treatment condition (N=3 donors). Statistical significance was determined by two-way ANOVA with Tukey's post-hoc test for multiple comparisons.

(H) Cellular proliferation is compared in HIV-1 infected, and uninfected bystander primary CD4+ T cells treated with DMSO, FASN-inhibitor or FASN-inhibitor with exogenous palmitate/oleate-BSA. Proliferation is determined by the retention of Cell Trace Violet. Statistical significance is determined by two-way ANOVA with Tukey's post-hoc test for multiple comparisons.

(I) Overview of fatty acid uptake assay and the expected outcome of experimental conditions (see methods). Primary CD4+ T cells are treated with palmitate-alkyne (50uM) for 30 minutes of exposure at either 37C, which facilitates uptake, or 4C, which inhibits uptake. Palmitate-BSA, which is not compatible with click chemistry, is utilized at matched concentration as a negative control. Alexa Fluor 647-Azide is utilized to label cells with palmitate alkyne uptake for detection by flow cytometry.

(J) Palmitate alkyne uptake is measured following click chemistry to assess the capacity of primary CD4+ T cells to internalize exogenous fatty acids. DMSO controls were compared to cells that were pretreated with FASN-inhibitor (GSK2194069) or FASN-inhibitor with exogenous palmitate-BSA and oleate-BSA. N=3 primary donors, and statistical significance was determined by two-way ANOVA with Tukey's post-hoc test for multiple comparisons.

*Statistical Significance:* *$p<0.05$, **$p\leq0.01$, ***$p\leq0.001$, ****$p\leq0.0001$.

*Abbreviations:* ACC = Acetyl-coA Carboxylase; FASN = Fatty acid synthase; DMSO = dimethyl sulfoxide; BSA = Bovine serum albumin (fatty acid free); FASNinh= FASN inhibitor; CTV = CellTrace Violet PA-Alkyne = palmitic acid alkyne; AF647 = Alexa Fluor 647.

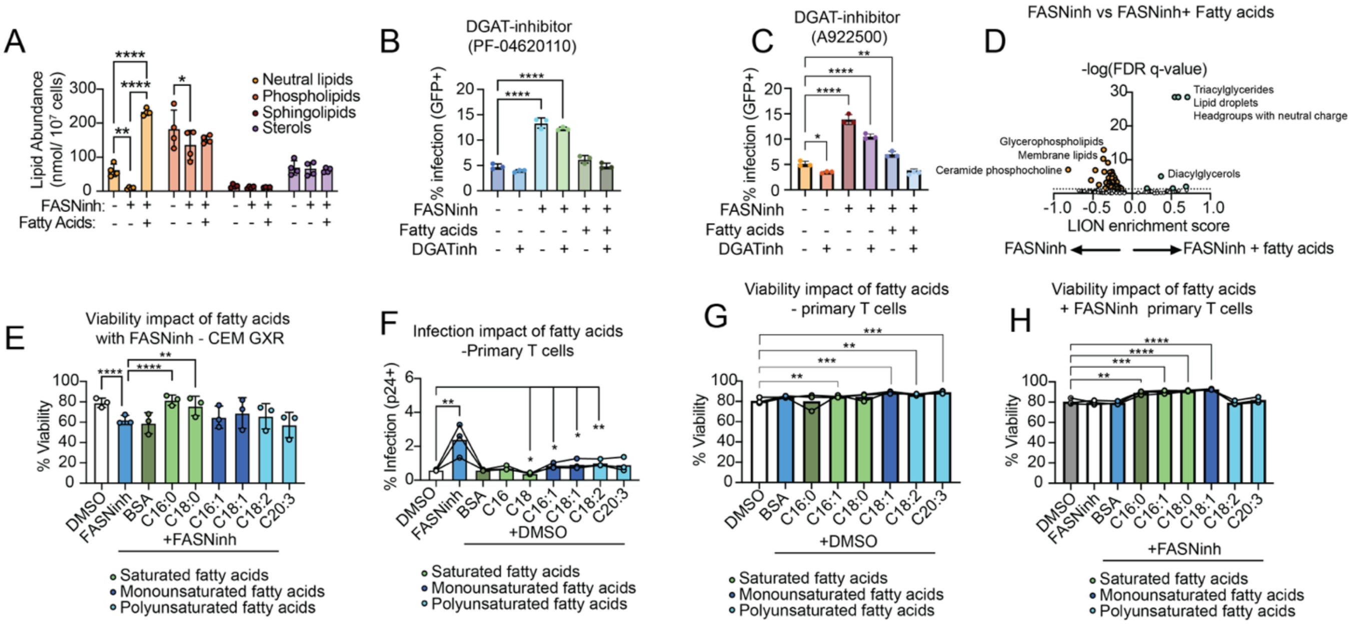


**Figure S3 (Related to Figure 3): Lipidomic analysis of FASN-inhibited T cells.**

(A) Impact of FASN inhibition with or without exogenous fatty acids on major lipid classes. Statistical significance was determined by two-way ANOVA with Tukey's multiple comparison test.

(B-C) The impact of DGAT1 inhibition is assessed on baseline infectivity, on cells under FASN inhibition, or cells under FASN inhibition with exogenous fatty acid exposure. Each datapoint represents a technical replicate (N=3), and statistical significance was determined by two-way ANOVA with Dunnet's test for multiple comparisons against the DMSO-treated controls. Two DGAT1 inhibitors were assessed, with PF-04620110 shown in B and A922500 shown in C.

(D) Enrichment analysis of hierarchical lipid-ontologies in FASN-inhibited cells with exogenous palmitate-BSA and oleate-BSA as compared to FASN-inhibited cells, as calculated utilizing the LION-web based ontology for lipid pathway enrichment. Hits that maintained an FDR adjusted q-value of ≤ 0.05 were indicated with orange (FASN-inhibited) or blue (FASNinh + fatty acids). For lipidomics analyses, N=4 technical replicates per group, as in Figure 3.

(E) The impact of exogenous fatty acids of increasing length and desaturation on the viability of CEM-GXR cells is assessed (N=3 experimental replicates). Statistical significance is calculated by two-way ANOVA with Dunnet's multiple comparison test against the FASN-inhibited condition.

(F) The impact of exogenous fatty acids of increasing length and desaturation on HIV-1 infectivity in DMSO-treated controls was assessed (N=3 primary donors). Statistical significance is calculated by two-way ANOVA with Dunnet's multiple comparison test against the FASN-inhibited condition.

(G-H) The impact of exogenous fatty acids of increasing length and desaturation on the viability of activated primary CD4+ T cells is assessed in cells that were pretreated with DMSO (G) or FASN-inhibitor (GSK2194069; H). Statistical significance is calculated by two-way ANOVA with Dunnet's multiple comparison test against the DMSO only condition (N=3 primary donors).

*Statistical Significance:* *$p<0.05$, **$p\leq0.01$, ***$p\leq0.001$, ****$p\leq0.0001$.

*Abbreviations:* nmol = nanomoles; DGATinh = Diacylglycerol O-Acyltransferase 1 Inhibitor; LION = Lipid Ontology; FASNinh = FASN inhibitor; FASN = Fatty acid Synthase; *DMSO* = dimethyl sulfoxide.

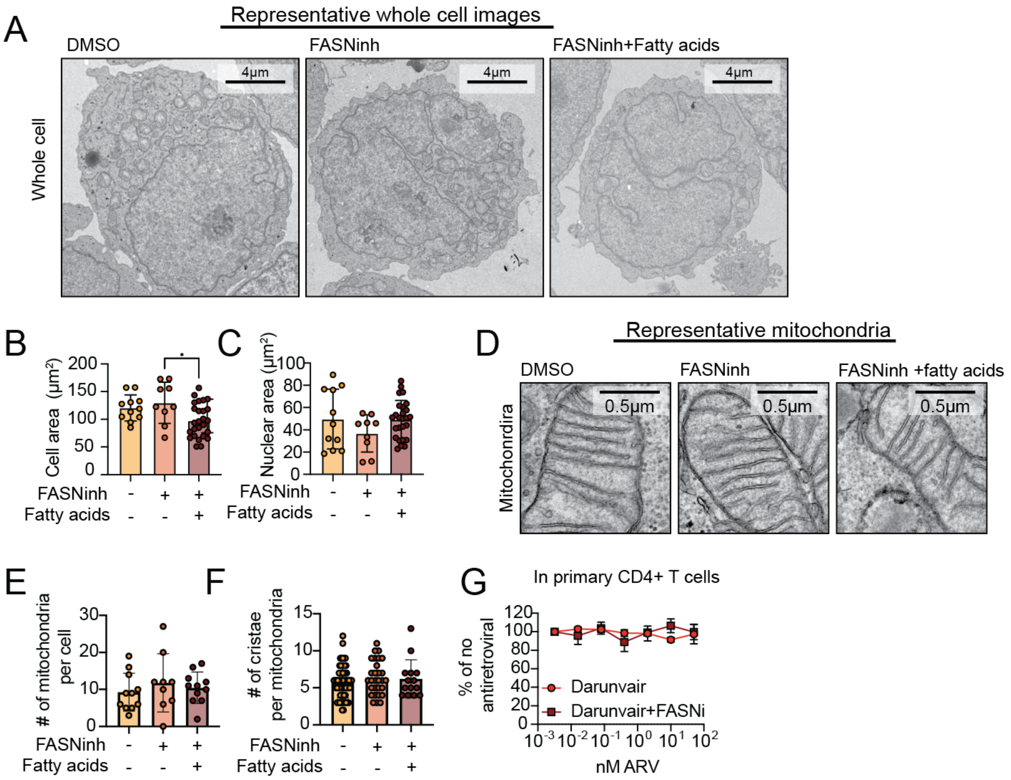


**Figure S4 (Related to Figure 4): Impact of FASN inhibition on cellular and nuclear architectures.**

(A) Representative whole-cell transmission electron microscopy (TEM) images are shown at 1400x magnification across treatment group. Scalebars represent 4 µm.

(B) Quantification of total cell area from 2800x images is compared across treatment group. Statistical significance is determined by one-way ANOVA with Tukey's post-hoc test for multiple comparisons.

(C) As with (B), but for the area of the nucleus.

(D) Representative TEM of mitochondria in each condition (magnification = 11,000x). Scalebars denote 500nm distances.

(E) The number of mitochondria per cell is compared across treatment group. Statistical significance is determined by one-way ANOVA with Tukey's post-hoc test for multiple comparisons.

(F) As with (E), but for the total number of visible cristae in each mitochondrion.

(G) The efficacy of Darunavir, which inhibits HIV-1 protease, on infectivity of HIV-1 pseudovirus in primary human T cells is compared with/without FASN inhibition. Each datapoint represents the average of 3 experimental replicates. Data could not be fit by non-linear 4-parameter regression.

*Statistical Significance:* *$p<0.05$, **$p\leq0.01$, ***$p\leq0.001$, ****$p\leq0.0001$.

*Abbreviations: FASN = fatty acid synthase; FASNinh = FASN inhibitor; DMSO =* dimethyl sulfoxide; ARV = antiretroviral
